# Unprecedented yet gradual nature of first millennium CE intercontinental crop plant dispersal revealed in ancient Negev desert refuse

**DOI:** 10.1101/2022.12.01.518650

**Authors:** Daniel Fuks, Yoel Melamed, Dafna Langgut, Tali Erickson-Gini, Yotam Tepper, Guy Bar-Oz, Ehud Weiss

## Abstract

Global agro-biodiversity has resulted from processes of plant migration and agricultural adoption. Although critically affecting current diversity, crop diffusion from antiquity to the middle-ages is poorly researched, overshadowed by studies on that of prehistoric periods. A new archaeobotanical dataset from three Negev Highland desert sites demonstrates the first millennium CE’s significance for long-term agricultural change in southwest Asia. This enables evaluation of the “Islamic Green Revolution” (IGR) thesis compared to “Roman Agricultural Diffusion” (RAD), and both versus crop diffusion since the Neolithic. Among the finds, some of the earliest *Solanum melongena* seeds in the Levant represent the proposed IGR. Several other identified economic plants, including two unprecedented in Levantine archaeobotany (*Ziziphus jujuba, Lupinus albus*), implicate RAD as the greater force for crop migrations. Altogether the evidence supports a gradualist model for Holocene-wide crop diffusion, within which the first millennium CE contributed more to global agro-diversity than any earlier period.

## Introduction

Crop diversity has long been recognized as key to sustainable agriculture and global food security, encompassing genetic resources for agricultural crop improvement geared at improving yields, pest resistance, climate change resilience, and the promotion of cultural heritage. Global genetic diversity of agricultural crops is a product of their dispersal from multiple regions and much research has attempted to reconstruct these trajectories [1-3]. As part of this effort, archaeobotanical research on plant migrations across the Eurasian continent has been a central theme in recent decades, especially with reference to “food globalization” and the “Trans-Eurasian exchange” [4-8]. Yet, as is true for archaeology-based domestication research in general, most studies of crop dispersal and exchange have focused on prehistoric origins and developments, to the near exclusion of more recent crop histories directly affecting today’s agricultural diversity [9-15]. One of the most influential, and contested, chapters in the later history of crop diffusion is the ‘Islamic Green Revolution’ (IGR) [16,17]. According to Andrew Watson, the IGR involved a package of sub-/tropical, mostly east- and south Asian domesticates which, as a result of the Islamic conquests, spread into Mediterranean lands along with requisite irrigation technologies ca. 700–1100 CE. This allegedly involved some 17 domesticated plant taxa (**Supplementary Table 1**), including such economically significant crops as sugar cane, orange and banana [16]. However, critics have argued that many of the proposed IGR crops were, and still are, of minor economic significance, while others were previously cultivated in the Mediterranean region, particularly under Roman rule, or else arrived much later [17-19]. Indeed, there is considerable evidence for crop diffusion immediately preceding and during the Roman period in the eastern Mediterranean, 1^st^ c. BCE– 4^th^ c. CE. During this time, several east- and central Asian crops, including some of those on Watson’s IGR list, were introduced to the Mediterranean region, along with agricultural technologies [17-21]. From this period on, a growing fruit basket is evident in sites and texts of the eastern Mediterranean region [22-25]. These include several tree-fruits (**Supplementary Table 2**) apparently reflecting the Greco-Roman passion for grafting and its pivotal role in the dispersal of temperate fruit crops from Central Asia to the Mediterranean and Europe [3,26]. Yet Roman arboricultural diffusion is but a subset of Roman agricultural diffusion (hereafter, RAD), which also includes non-arboreal crops (including cannabis, muskmelon, white lupine, rice, sorghum) and various agricultural techniques diffused by the Romans into the eastern Mediterranean [21,27-35]. Not all crops in motion during this period took hold in local agriculture. In some cases, as has been claimed for rice in Egypt, initial Roman-period importation of the new crops ultimately led to local cultivation in the Islamic period [36]. In other cases, Roman introductions were subsequently abandoned [37], or failed to diffuse beyond elite gardens until much later [38]. Limited adoption in local agriculture is also a feature of some proposed IGR crops, as Watson admitted regarding coconut and mango [16]. Thus, a cursory consideration of proposed IGR and RAD crops in the eastern Mediterranean reveals that the balance between the two is about even and perhaps weighted toward RAD (**Supplementary Tables 1-2**). This sort of comparison is valuable for evaluating the IGR thesis and attaining improved understandings of crop exchange and dispersal in the first millennium CE, but a higher-resolution micro-regional approach is needed to rigorously gauge these developments. Systematic evaluation of relative Islamic and Roman contributions to agricultural dispersal has been attempted for Iberia [35,39]. In the eastern Mediterranean, archaeobotanical studies in Egypt [36], northern Syria [40], and Jerusalem [25,41-42] have also yielded evidence for IGR introductions framed against Roman agricultural diffusion, but these have not yet been considered holistically.

The exceedingly rich plant remains from relatively undisturbed Negev Highland middens (**Fig. 1-2;** [43-45]) provide a significant new addition to the evidence for Levantine and Mediterranean crop diffusion, informing upon changes in the local economic plant basket over the 1^st^ millennium CE. The Negev Highlands also offer an ideal test case for the geographical extent of crop dispersal, as a desert region on the margins of the settled zone, which practiced vibrant runoff farming and engaged in Mediterranean and Red Sea trade networks of Late Antiquity [46-50]. Archaeobotanical finds from the Negev Highlands, mainly from Byzantine sites (5^th^-7^th^ centuries CE), have been reported in previous studies [43-44,51-59], including those deriving from organically rich middens at Elusa, Shivta, and Nessana, excavated as part of the recent NEGEVBYZ project [53-59]. We present below the first complete dataset of identified plant remains from the Late Antique Negev Highland middens dated to the local Roman, Byzantine and early Islamic periods (2nd-8^th^ centuries CE). We then analyze this data to assess the evidence for Roman and Early Islamic crop diffusion in the southern Levant, comparing with earlier introductions. These include the southwest Asian Neolithic ‘founder crops’, Chalcolithic-Early Bronze Age tree fruit domesticates, and Bronze-Iron Age introductions (**Supplementary Tables 1-3**). This analysis offers Holocene-scale insights on the dynamics of crop diffusion.

**Figure 1.**
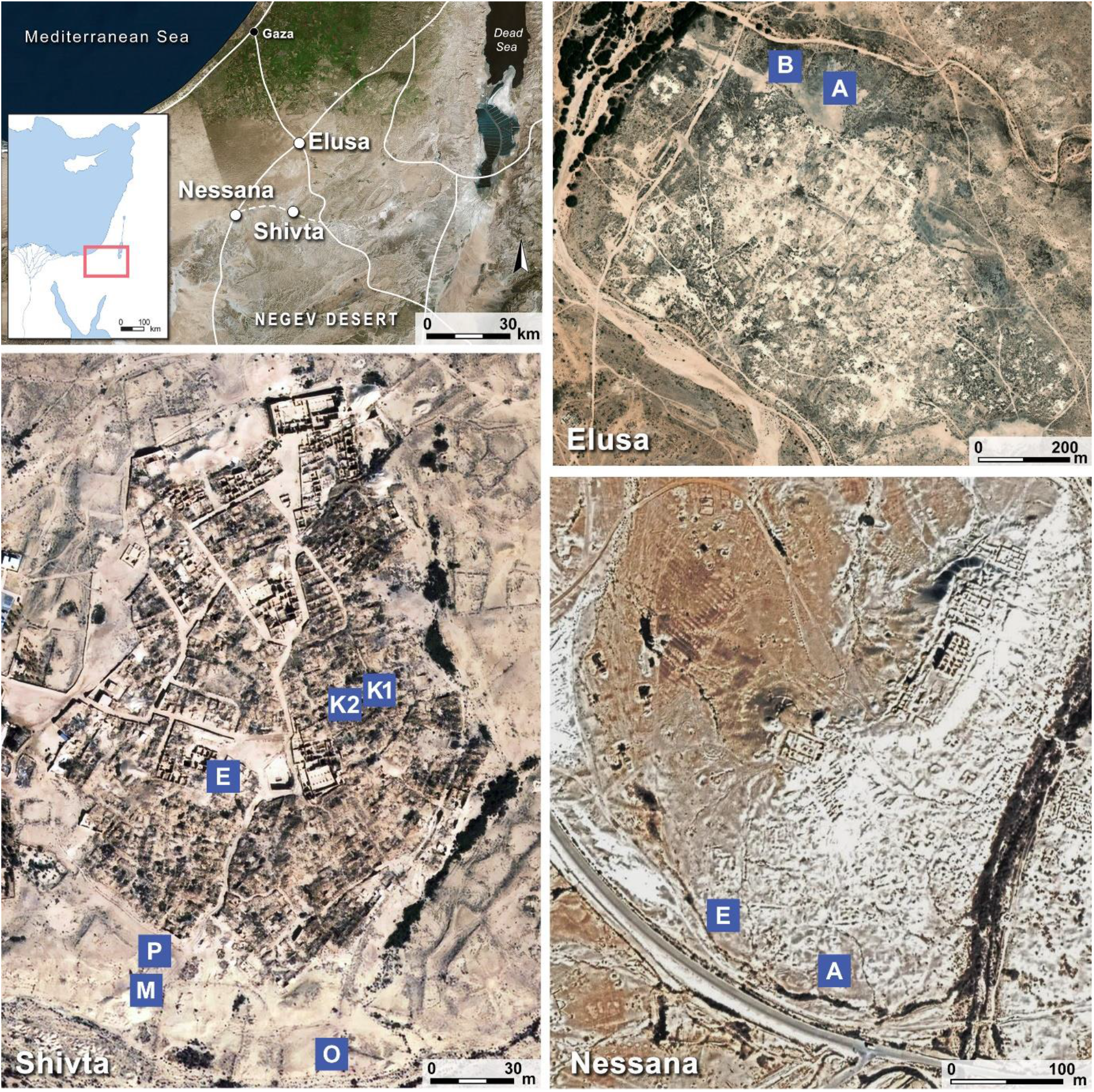
Study sites and middens The study sites – Shivta, Elusa and Nessana – roughly span the Negev Highlands region of the Negev desert. The excavated middens are marked on the aerial photos above. Middens are lettered as named in the 2015-2017 excavations (see also Table 2).

## Results

Roughly 50,000 quantifiable macroscopic plant parts were retrieved from fine-sifted flotation and dry-sieved sediment samples of the middens of Elusa, Shivta and Nessana, excluding charcoal and in addition to a roughly equal number retrieved from wet-sieving (see **Supplementary Information**). These mostly carpological remains were identified to a total 144 distinct plant taxa (**Supplementary Table 4**). Nearly half of the identified specimens derived from six Shivta middens; one quarter from three Elusa middens and one quarter from two Nessana middens. Preservation quality varied somewhat within and between middens and samples, but all middens yielded rich concentrations of charred seeds and other organic remains, including many exceptionally preserved specimens. Identified species were classified as either domestic or wild and the former grouped by functional category (**Supplementary Table 4**). Most of the 120 wild taxa have ethnographically documented uses, whether for forage or fodder, crafts or fuel, food or spice, medicine or recreation.

Nearly all of them grow wild in the Negev Highlands today and we cannot determine for certain which were deliberately used on site. Twenty-three domesticated food plant types were identified, including cereals, legumes, fruits, nuts, and one vegetable. Like the other domesticates, we consider the presence of Nile acacia (*Vachellia nilotica* [L.] Willd. ex Delile) in the assemblage to be the result of deliberate import or cultivation, along with other exotic trees previously identified by charcoal and pollen from the study sites. We focus on these 24 plants as indicators of local foodways and global crop diffusion. Their presence/absence by period in the Negev Highland middens appears in **Table 1**, and orders of magnitude by midden context for fine-sifted archaeobotanical samples appear in **Table 2** (see **Supplementary Information** for sifting and sampling strategy). The latter enable categorization of the Late Antique Negev Highland domesticates as staples, cash crops, and luxury/supplementary foods, setting the stage for analysis of the local manifestation of long-term crop diffusion. This analysis is further augmented by identified charcoal and pollen data from the study sites (**Supplementary Tables 5-6**) which raise the number of distinct plant taxa identified in the NEGEVBYZ project to over 180. Among the charcoal/pollen taxa not identified by seed and fruit remains are three fruit trees: sycomore fig (*Ficus sycomorus* L.), doum palm (*Hyphaene thebaica* [L.] Mart.), and hazelnut (*Corylus* sp.).

**Table 1.**
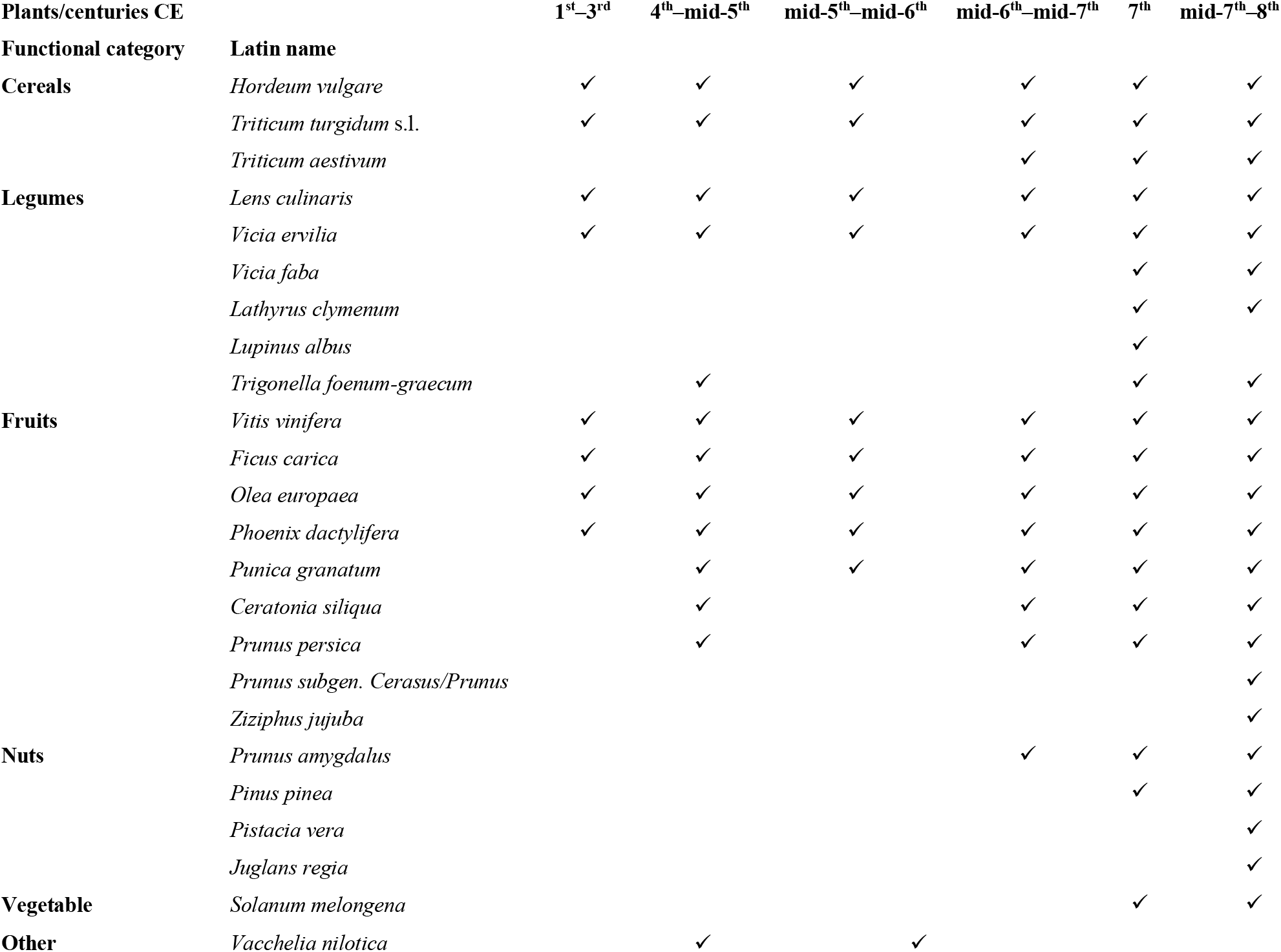
Presence/absence of domesticated species in Negev Highland middens by period (carpological remains)

Seed quantities and ubiquity point to barley (*Hordeum vulgare* L.), wheat (*Triticum turgidum/aestivum*), and grapes (*Vitis vinifera* L.) as the main cultivated crops, which were clearly calorific staples. Their local cultivation is attested to by cereal processing waste (rachis fragments, awn and glume fragments, culm nodes and rhizomes) and wine-pressing waste (grape pips, skins, and pedicels). In addition, lentil (*Lens culinaris* [L.] Coss. & Germ.), bitter vetch (*Vicia ervilia* [L.] Willd.), fig (*Ficus carica* L.), date (*Phoenix dactylifera* L.), and olive (*Olea europaea* L.) should also be counted as staples based on seed quantities and ubiquity (**Tables 1-2**). They were likely cultivated locally. Significantly, all identified staples were among the southwest Asian Neolithic founder crops and early fruit domesticates which formed a stable part of Levantine diets by the Early Bronze Age (3300–2000 BCE).

**Table 2.** Domesticated plant seeds order of magnitude by period, site, and area (from fine-sift)

| Century CE | 1<sup>st</sup>–<br>3<sup>rd</sup> | 4<sup>th</sup>–<br>mid-5<sup>th</sup> | mid-5<sup>th</sup>–<br>mid-6<sup>th</sup> |  | mid-5<sup>th</sup>–<br>mid-7<sup>th</sup> | mid-6<sup>th</sup>–<br>mid-7<sup>th</sup> |  | early 7<sup>th</sup> |  | 7<sup>th</sup>–<br>8<sup>th</sup> | mid-7<sup>th</sup>–8<sup>th</sup> |  |  |  |
| --- | --- | --- | --- | --- | --- | --- | --- | --- | --- | --- | --- | --- | --- | --- |
| Site | SVT | HLZ | HLZ | SVT | NZN | NZN | SVT | SVT |  | NZN | NZN | SVT |  |  |
| Area (midden) | P | A4 | A1 | M | A | A | O | K2 | E | A | E | K1 | K2 | E |
| Samples | 5 | 14 | 19 | 14 | 7 | 5 | 12 | 3 | 3 | 27 | 10 | 13 | 13 | 12 |
| Vol. (L) | 15 | 85 | 85 | 42 | 21 | 15 | 36 | 9 | 9 | 84 | 33 | 39 | 39 | 36 |
| Plant species |  |  |  |  |  |  |  |  |  |  |  |  |  |  |
| *Hordeum vulgare* | XX | XXX | XXX | XX | XXX | XX | XX | XXX | XXX | XXX | XXX | XXX | XXX | XXX |
| *Triticum* sp. | XX | XX | XX | XX | X | X | X | XX | XX | XX | XXX | XXX | XXX | XXX |
| *Lens culinaris* |  | XX | XX | X | XX |  | X | X | X | X | XX | XX | X | X |
| *Vicia ervilia* | X | X | X | X | X | X | X | X | XX | X | XX | XX | X | XX |
| *Trigonella foenum-graecum* |  | X |  |  |  |  |  |  | X | X | X | X | X |  |
| *Lathyrus clymenum* |  |  |  |  |  |  |  |  |  | X |  | X |  |  |
| *Lupinus albus* |  |  |  |  |  |  |  |  |  |  |  | X |  |  |
| *Vitis vinifera* | X | XX | XX | XX | XX | X | XX | XX | X | XXX | XXX | XXX | XXX | XX |
| *Ficus carica* | X | XXX | XXX | XX | X | X | XX | X | X | XX | X | X | XX |  |
| *Olea europaea* |  | X |  | X | X | X | X | X |  | X | XX | X | X | X |
| *Phoenix dactylifera* | X | X | X | X | X |  | X | X | X | X | XX | XX | X | X |
| *Punica granatum* |  | rind |  | rind | X | rind | X | rind |  | X | XX | X | X | X |
| *Ceratonia siliqua* |  |  |  |  |  |  |  |  |  | X |  | X | pistil |  |
| *Prunus amygdalus* |  |  |  |  |  |  |  |  |  | X |  | X | X | X |
| *Prunus persica* |  | X |  |  |  |  | X |  |  | X |  | X |  |  |
| *Pinus pinea* |  |  |  |  |  |  |  |  |  | X | X |  |  |  |
| *Solanum melongena* |  |  |  |  |  |  |  |  |  | X |  |  |  | X |
| *Vachellia nilotica* |  | X | X |  | X |  |  |  |  |  |  |  |  |  |

Grapes were previously shown to be the primary cash crop of the Byzantine Negev Highlands—particularly in the mid-5^th^ to mid-6^th^ c. CE—based on their changing relative frequencies [54]. Yet, we cannot rule out the possibility of cereal cultivation for export in some periods. One modern example is the export of Negev barley to Britain for beer production in the 19^th^ century [60]. Interestingly, free-threshing hexaploid bread wheat (*Triticum aestivum* L.)—a more market-oriented wheat species identifiable archaeologically by indicative rachis segments—appears in the Negev Highlands only after the mid-6^th^ c. (**Table 2**). This corresponds with the period of decline in viticulture [54].

In the ‘luxuries and supplements’ category we include potentially important and desirable dietary components which were minor and apparently nonessential in local consumption or agriculture. These include several food crops poorly represented in the local assemblages: fava bean (*Vicia faba* L.), fenugreek (*Trigonella foenum-graecum* L.), Spanish vetchling (*Lathyrus clymenum* L.), and white lupine (*Lupinus albus* L.) among the legumes; peach (*Prunus persica* [L.] Batsch), plum/cherry (*Prunus* subgen. *Cerasus/Prunus*), carob (*Ceratonia siliqua* L.) and jujuba (*Ziziphus jujuba* Mill.) among the tree-fruits; almond (*Prunus amygdalus* Batsch), walnut (*Juglans regia* L.), stone pine (*Pinus pinea* L.), pistachio nut (*Pistacia vera* L.) and hazel (*Corylus* sp.) among the nuts; the aubergine (*Solanum melongena* L.) as a unique summer vegetable (**Fig. 2-3**); and supplementary wild edibles such as beet (*Beta vulgaris* L.), coriander (*Coriandrum sativum* L.), and European bishop (*Bifora testiculata* [L.] Spreng.) (**Supplementary Table 4**). Any of these could have been cultivated in Negev Highland runoff farming [47, 59], or on site [61].

**Figure 2.**
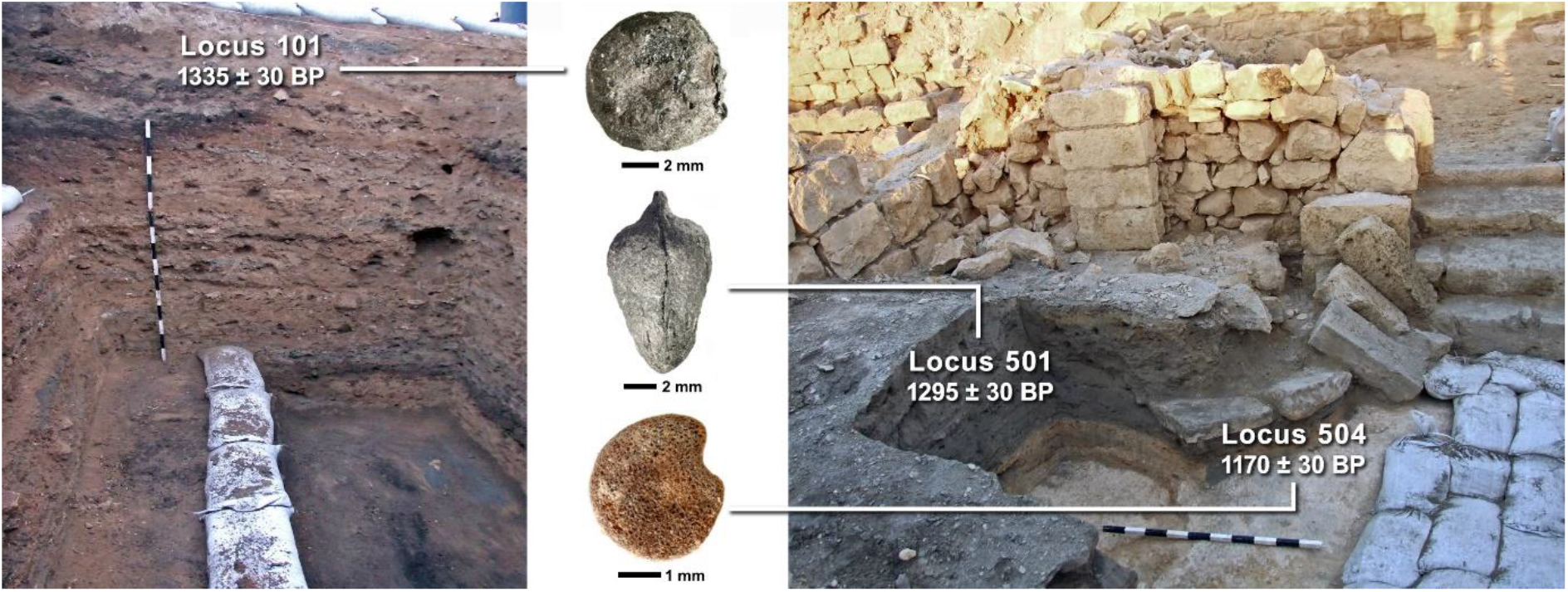
First finds from the Negev Highlands middens *Section photos of Nessana midden A (left) and Shivta midden E (right) are shown with select Loci and their uncalibrated radiocarbon dates (photographed by: Yotam Tepper), from which seeds of* Lupinus albus *(center top)*, Ziziphus jujuba *(center middle)*, Solanum melongena *(center bottom) were found. These seeds represent some of the earliest of their species found in the southern Levant (photographed by Daniel Fuks).*

Another important ancient economic plant found in the assemblages is the Nile acacia, which does not grow today in the Negev. Previous archaeobotanical finds of Nile acacia in the Levant all come from Roman-period sites in the Dead Sea rift valley, which Kislev [62] interpreted as a component of the ancient flora in this region of Sudanian vegetation penetration. However, this was also an important region for desert-crossing camel caravan commerce. Nile acacia seed finds from Elusa (**Fig. 3**) are the first from outside the phytogeographic region of Sudanian vegetation, but they remain within the ancient caravan trade routes connecting the Red Sea and the Mediterranean. Therefore, we consider Nile acacia seeds to represent a Roman-period introduction to the Levant, whether as objects of cultivation or of trade at the Negev desert route sites. Other exotic trees used for quality wood and craft were identified by pollen and/or charcoal, including: cedar of Lebanon (*Cedrus libani* A.Rich.), European ash (*Fraxinus excelsior* L.), and boxwood (*Buxus sempervirens* L.). Cedar was identified by both charcoal and pollen, suggesting local garden cultivation (see Langgut et al. 2021 [59] and **Table 3**).

**Table 3.** Combined evidence for fruit/nut trees

| Taxon | Common name | Seeds/Fruit |  |  | Charcoal |  |  | Pollen |  |  |
| --- | --- | --- | --- | --- | --- | --- | --- | --- | --- | --- |
|  |  | SVT | NZN | HLZ | SVT | NZN | HLZ | SVT 1 | SVT 2 | SVT 3 |
| *Vitis vinifera* | grapevine | + | + | + | + | + | - | + | + | + |
| *Olea europaea* | olive | + | + | + | + | - | + | + | + | + |
| *Ficus carica* | common fig | + | + | + | + | + | + | - | - | - |
| *Phoenix dactylifera* | date palm | + | + | + | + | - | - | + | + | + |
| *Ceratonia siliqua* | carob | + | + | + | - | - | - | + | - | + |
| *Punica granatum* | pomegranate | + | + | + | - | + | - | - | - | - |
| *Prunus* spp. | almond/peach/plum | + | + | + | + | + | - | - | - | - |
| *Pinus* spp. | pine | + | + | - | + | + | + | + | + | + |
| *Corylus* sp. | hazel | - | - | - | - | - | - | + | - | + |
| *Ficus sycomorus* | sycomore fig | - | - | - | - | + | + | - | - | - |
| *Hyphaene thebaica* | doum palm | - | - | - | + | + | - | - | - | - |
| *Juglans regia* | walnut | + | - | - | - | - | - | - | - | - |
| *Pistacia vera* | pistachio | + | - | - | * | * | - | - | - | - |
| *Ziziphus jujuba* | jujuba | + | - | - | * | * | - | - | - | - |

*Carpological, anthracological and palynological evidence for fruit- and exotic trees in the study sites. Assessment of local cultivation is based on the combination of proxies and especially pollen to include grapevine, fig, olive, date, pomegranate, carob, hazelnut, cedar and the Prunus genus (potentially including almond, peach, plum and/or cherry). Local cultivation of stone pine may also plausibly be inferred. SVT1= South reservoir, Shivta; SVT 2 = North reservoir, Shivta; SVT3 = North church garden, Shivta; + indicates presence; - indicates absence; \*indicates charcoal identified to genus, including possible local wild species. Prunus spp. includes Prunus dulcis and Prunus domestica/cerasus endocarp/exocarp, and Prunus spp. charcoal. Pinus spp. includes Pinus pinea seed coats, Pinus halepensis charcoal, and Pinus sp. pollen.*

**Figure 3.**
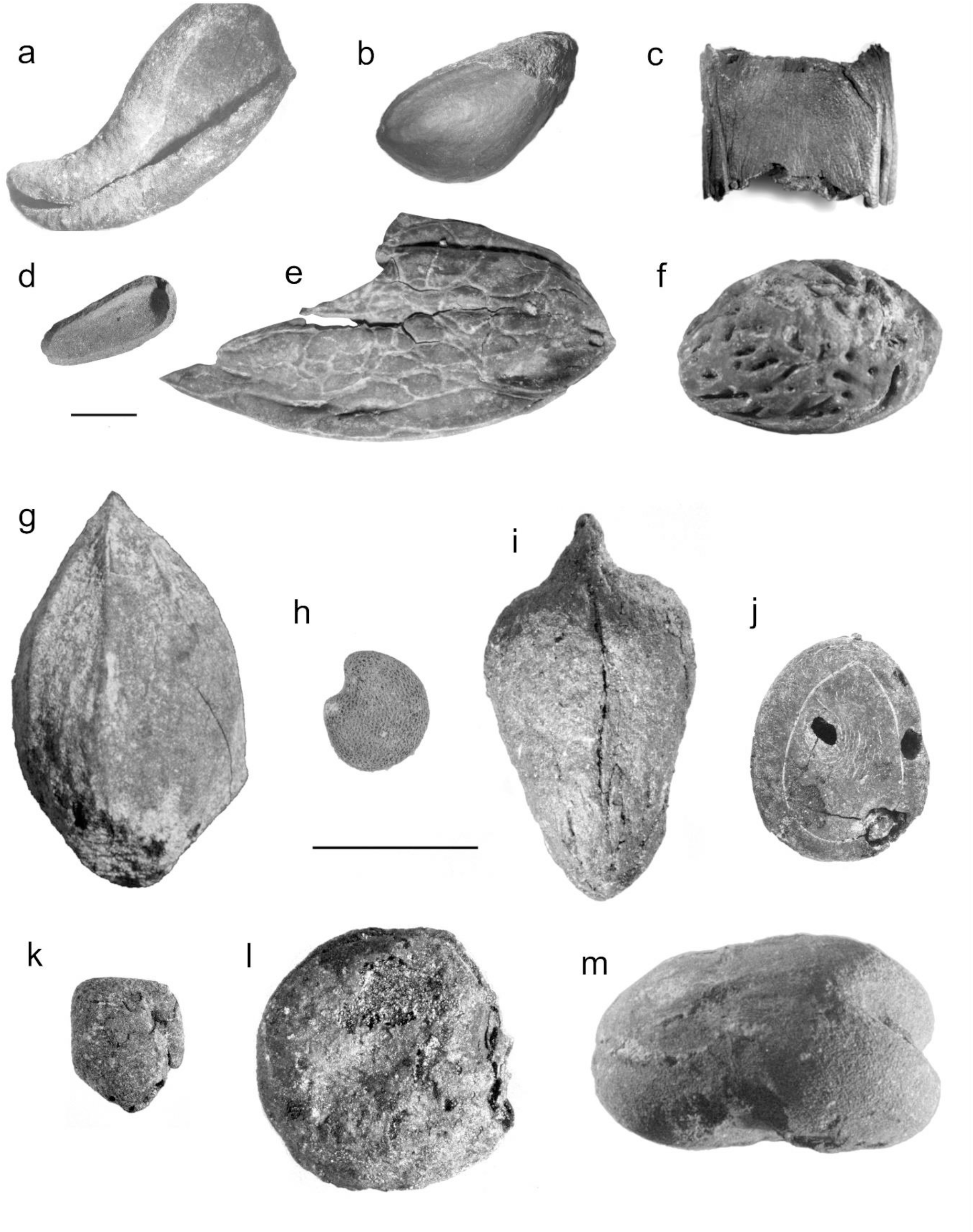
Select plant remains from the Negev Highland middens *(a) charred almond* (Prunus amygdalus *Batsch.) exocarp; (b) charred pistachio (*Pistacia vera *L.) drupe; (c) charred carob (*Ceratonia siliqua *L.) pod fragment; (d) uncharred stone pine (*Pinus pinea *L.) outer seed coat fragment; (e) uncharred walnut (*Juglans regia *L.) endocarp of the thin-shelled variety (f) charred peach (*Prunus persica *[L.] Batsch) endocarp; (g) charred cherry/plum (*Prunus *subgen.* Cerasus/Prunus) *endocarp; (h) uncharred aubergine (*Solanum melongena *L.) seed; (i) charred jujuba (*Ziziphus jujuba *Mill.) endocarp; (j) charred* Vachellia nilotica *(L.) P.J.H.Hurter & Mabb. seed; (k) charred fenugreek (*Trigonella foenum-graecum/berythea*) seed; (l) charred white lupine (*Lupinus albus *L.) seed; (m) charred fava bean (*Vicia faba *L.). Scale bars = 5mm; all photos in grayscale (photographed by: Daniel Fuks and Yoel Melamed)*.

Complementing the seed/fruit remains presented above, palynological and anthracological analyses support local cultivation of grapevine, fig, olive, date, pomegranates, carob, and the *Prunus* genus, which includes almond, peach, plum and/or cherry [59]. Based on stone pine seed coats, and the identification of Pinaceae pollen (= pine other than the local Aleppo pine), it is plausible that stone pine was cultivated locally, albeit on a small scale (**Table 3**). Pollen evidence also supports local cultivation of hazel – another domesticate unattested in the southern Levant before the Roman period **(Tables 3, 5; Supplementary Tables 5-6**).

Overall, the later-period middens were more concentrated in plant remains, and it is in the Early Islamic period middens where we find most of the rare domesticated species, RAD crops included (**Table 1**). This appears to be related to taphonomy, and therefore absence of RAD crops in the Byzantine middens should not be taken as evidence of their absence (see **Supplementary Information**). Samples containing the unique finds of white lupine and jujuba – which are unprecedented in southern Levantine archaeobotany – were dated to the Umayyad or early Abbasid period (mid-7^th^ – late 8^th^ c. cal. CE at 2σ; see **Fig. 1**; **Table 4** and **Supplementary Information**). However, textual studies have identified these species in Roman-period texts of the southern Levant [22]. The sample from Shivta containing aubergine seeds was dated to the Abbasid period (772-974 cal CE at 2σ), supporting previous finds from Abbasid Jerusalem [25,40-41].

**Table 4.** Radiocarbon dating of select loci

| Radiocarbon Lab. no. | Site | Locus | Basket | Special find | Material dated | Uncal BP | Cal. CE (1 $\sigma$) | Cal. CE (2 $\sigma$) |
| --- | --- | --- | --- | --- | --- | --- | --- | --- |
| Poz-141223 | Nessana | 101 | 1040-1 | white lupine | charred barley seed | 1335 ± 30 | 654 (46.4%) 682 | 647 (61.0%) 708 |
|  |  |  |  |  |  |  | 745 (18.3%) 760 | 730 (34.5%) 775 |
|  |  |  |  |  |  |  | 768 ( 3.6%) 771 |  |
| Poz-141225 | Shivta | 504 | 5029 | aubergine | charred barley seed | 1170 ± 30 | 776 ( 9.7%) 788 | 772 (73.9%) 901 |
|  |  |  |  |  |  |  | 825 (49.0%) 894 | 916 (21.6%) 974 |
|  |  |  |  |  |  |  | 928 ( 9.6%) 945 |  |
| Poz-141226 | Shivta | 501 | 5108 | jujuba | charred barley seed | 1295 ± 30 | 670 (33.4%) 704 | 659 (95.4%) 775 |
|  |  |  |  |  |  |  | 739 (34.9%) 772 |  |

Considering together the domestic plants evident in the Negev Highlands according to their period of first attestation in the southern Levant – archaeobotanically and historically – offers a window onto processes of long-term crop diffusion (**Table 5)**. While their quantities and ubiquities indicate that RAD and IGR crops were initially of minor significance, they make up over a third of the domesticates’ species diversity (**Fig. 4; Table 5**). All the more surprising considering the Negev Highlands’ desert and present-day peripheral status, this new data reveals for the first time the extent of western influence on local agriculture and trade (**Fig. 5**).

**Table 5.** Earliest archaeobotanical evidence in the southern Levant for domestication/introduction of Negev Highland domesticated plants

| Latin name | Period | Tag | Approx date | Reference |
| --- | --- | --- | --- | --- |
| *Hordeum vulgare* | PPNB | Founder crop | 9<sup>th</sup> mill. BCE | Zohary et al. 2012 [3] |
| *Lens culinaris* | PPNB | Founder crop | 9<sup>th</sup> mill. BCE | Caracuta et al. 2017 [74] |
| *Vicia ervilia* | PPNB | Founder crop | 9<sup>th</sup> mill. BCE | Caracuta et al. 2017 [74] |
| *Vicia faba* | PPNB | Founder crop | 9<sup>th</sup> mill. BCE | Caracuta et al. 2017 [74] |
| *Triticum turgidum* s.l. (free-threshing) | PPNB | Founder crop | 7<sup>th</sup> mill. BCE | Feldman and Kislev 2007 [75] |
| *T. aestivum* (free-threshing) | NA | Founder crop | NA | Zohary et al. 2012 [3] |
| *Olea europaea* | Chalcolithic/E. Bronze | Early fruit domesticate | 5<sup>th</sup> mill. BCE | Langgut et al. 2019 [76] |
| *Ficus carica* | Chalcolithic/E. Bronze | Early fruit domesticate | 5<sup>th</sup> mill. BCE | Weiss 2015 [77] |
| *Vitis vinifera* | Chalcolithic/E. Bronze | Early fruit domesticate | 5<sup>th</sup> mill. BCE | Weiss 2015 [77] |
| *Phoenix dactylifera* | Chalcolithic/E. Bronze | Early fruit domesticate | 5<sup>th</sup> mill. BCE | Weiss 2015 [77] |
| *Punica granatum* | Chalcolithic/E. Bronze | Early fruit domesticate | 5<sup>th</sup> mill. BCE | Melamed 2002 [78] |
| *Prunus amygdalus* | Chalcolithic/E. Bronze | Early fruit domesticate | 5<sup>th</sup> mill. BCE | Zohary et al. 2012 [3] |
| *Lathyrus clymenum* | Middle Bronze | Bronze Age introduction | 19<sup>th</sup>-18<sup>th</sup> c. BCE | Kislev et al. 1993 [79] |
| *Juglans regia* | Middle Bronze | Bronze Age introduction | 18<sup>th</sup> c. BCE | Langgut 2015 [80] |
| *Trigonella foenum-graecum* | Late Bronze Age IIA | Bronze Age introduction | 14<sup>th</sup> c. BCE | Weiss et al. 2019 [81] |
| *Prunus persica* | Nabatean | RAD crop | 1<sup>st</sup> c. BCE | Kislev and Simchoni 2009 [82] |
| *Vachellia nilotica* | Nabatean | RAD crop | 1<sup>st</sup> c. BCE | Kislev 1990 [62] |
| *Ceratonia siliqua* | Hellenistic-Roman | RAD crop | 1<sup>st</sup> c. BCE | Zohary et al. 2012 [3] |
| *Pinus pinea* | Hellenistic-Roman | RAD crop | 1<sup>st</sup> c. BCE | Kislev 1988 [83] |
| *Prunus* subgen. *Cerasus*/*Prunus* | Roman | RAD crop | 1<sup>st</sup> c. CE | Tabak 2006 [84] |
| *Pistacia vera* | Roman | RAD crop | 2<sup>nd</sup> c. CE | Hartman and Kislev 1998 [85] |
| *Corylus* sp. | Roman | RAD crop | 2<sup>nd</sup> c. CE | Kislev and Simchoni 2006 [23];<br>Langgut et al. 2021 [59] |
| *Lupinus albus* | Early Islamic | RAD crop | 7<sup>th</sup> c. CE | this paper |
| *Ziziphus jujuba* | Early Islamic | RAD crop | 7<sup>th</sup> c. CE | this paper |
| *Solanum melongena* | Early Islamic | IGR crop | 7<sup>th</sup> c. CE | Amichay et al. 2019 [25]; this<br>paper |

*Note: The earliest evidence for Prunus subgen. Cerasus/Prunus refers to plum (Prunus subgen. Prunus) only. Cherry (Prunus* *subgen. Cerasus) has yet to be identified in the southern Levantine archaeobotanical record.*

**Figure 4.**
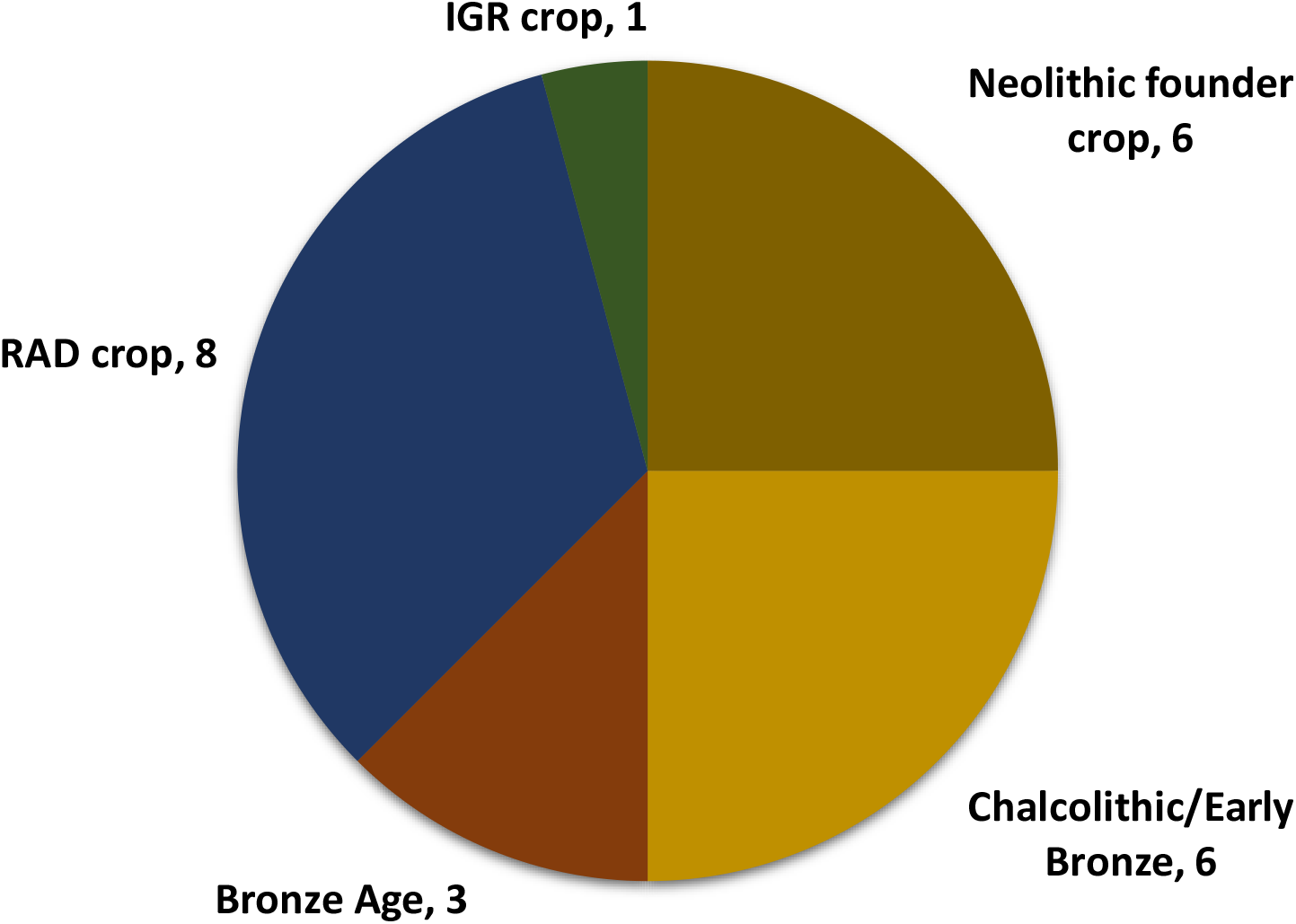
Negev Highlands crop basket by period of introduction to the southern Levant (based on carpological remains)

**Figure 5.**
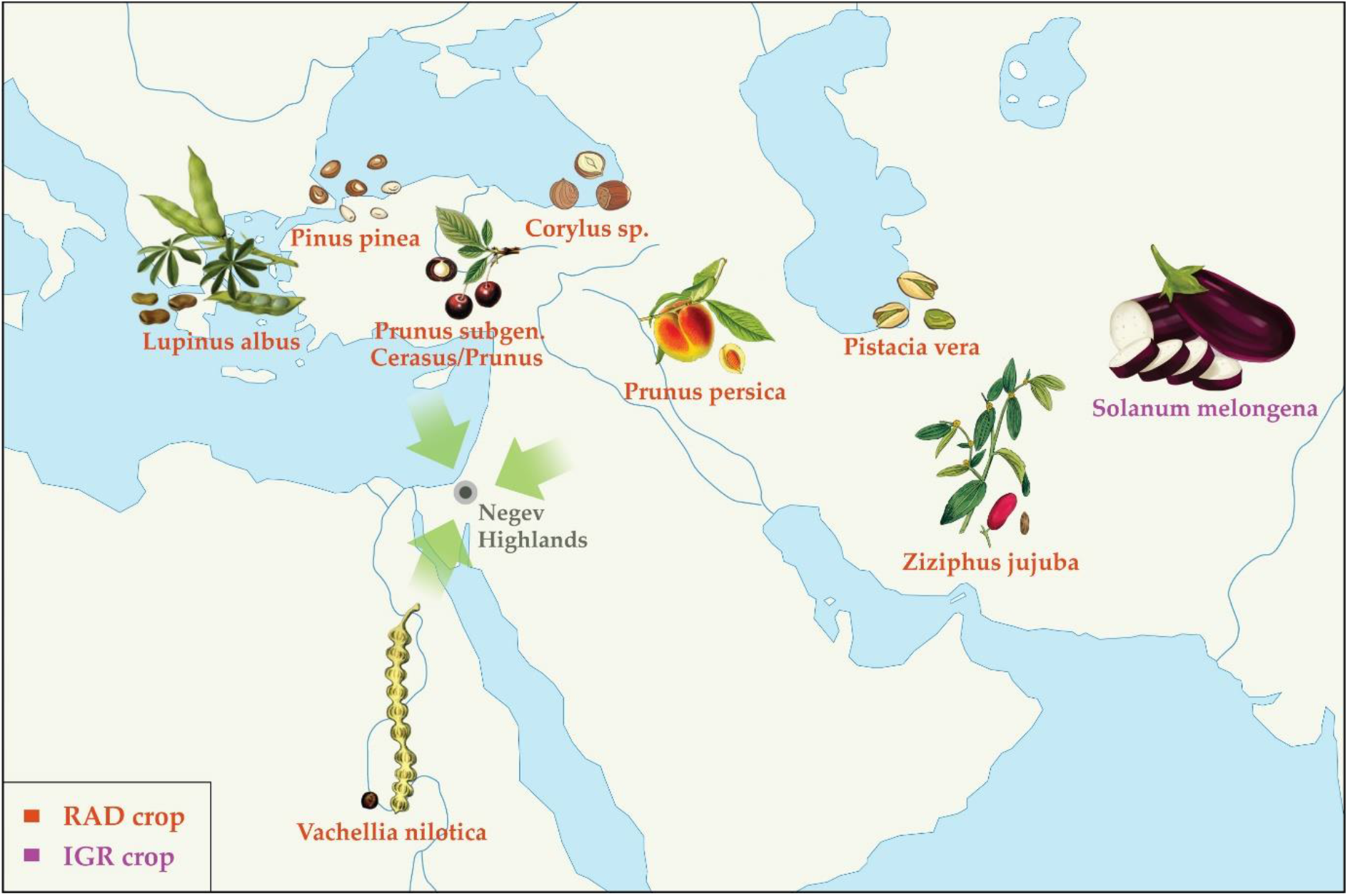
First mill. CE southern Levantine introductions found in Negev Highland middens Schematic representation of directions of first millennium CE crop diffusion into the southern Levant based on plants attested to in the Negev Highland middens. RAD crops are labeled red; IGR crops purple.

## Discussion

The critical mass afforded by the new, systematically retrieved and identified plant remains from Late Antique Negev Highland trash mounds allows not only reconstructions of local plant economy, but also insights on the dispersal of crop plants over the last 11.5 ky. Of the Negev Highland plant remains, only the aubergine is an IGR crop (**Table 5; Fig. 4-5**).

Together with finds from Abbasid Jerusalem, seeds found in the Negev Highland middens are among the earliest archaeobotanical finds of this plant in the Levant and are roughly contemporary with the earliest textual references to aubergine [16,22]. Significantly, aubergine is the only summer crop in the Negev Highlands plant assemblage. In other regions of the southern Levant, summer crops were certainly cultivated in the Roman period [20,63], but the Early Islamic introduction of aubergine is consistent with Watson’s claim that summer cultivation expanded in this later period [16,64]. Ultimately, widespread adoption of summer-winter crop rotation in the Mediterranean region effected changes in people’s diets and work routines. Yet these changes clearly did not occur overnight. To be fair, the Early Islamic assemblages from the Negev Highlands do not offer enough of a time perspective to fully gauge the effects of Early Islamic crop introduction on their own as they span only the first 200-300 years of Islam. Yet it is also possible that finds from the 7^th^-8^th^ century middens represent Byzantine agronomic traditions and techniques. Regardless, had crop introductions been inundating and pervasive during the Early Islamic period, we expect they would have been more apparent in Negev Highland crop diversity.

By contrast, the Negev Highlands crop basket highlights the influence of RAD, particularly on arboriculture. Of the 24 domestic plants identified by carpological remains, seven were introduced to the southern Levant during the 1^st^ c. BCE to the 4^th^ c. CE: pistachio nut, stone pine, peach, plum/cherry, jujuba, Nile acacia, white lupine, plus carob which is a local wild species but was apparently not fully domesticated until the Classical period (**Table 5**). Jujuba and white lupine are unprecedented in southern Levantine archaeobotany, but they are known from Roman-period texts and the archaeobotany of neighboring regions [65-68]. Considering pollen remains, hazelnut is an additional RAD species identified in the Negev Highlands, that was also found in Herod’s garden at Caesarea, probably as an imported ornamental [69]. The fact that the RAD plant remains are more prevalent in the Early Islamic phase (**Table 1-2**) is likely the result of overall better preservation and plant richness in this phase. Therefore, we understand them to be part of the general Late Antique Negev Highlands domestic plant assemblage, noting that their earliest secure archaeobotanical records in the southern Levant as a whole derive mostly from the 1^st^ c. BCE to the 2^nd^ c. CE (**Table 5**). We acknowledge that some RAD species are first attested to at the end of the Hellenistic period of the southern Levant in the 1^st^ c. BCE. We nonetheless consider them RAD crops in view of chronological proximity and their entrenchment in local agriculture and culture during the Roman period.

Allowing for gaps in the archaeobotanical record, partially compensated by textual references, it is still fair to say that the RAD plants—which comprise a significant proportion of species diversity in the Late Antique Negev Highland basket of domestic plants—were introduced to the southern Levant over a relatively short period in Holocene history.

The snapshot presented here of the Negev Highlands’ microregional crop basket supports and significantly enhances previous evidence for 1^st^ millennium CE crop diffusion. Together with the archaeobotany of sites from southern Jordan [70] and Jerusalem [25,41], the Negev Highland plant remains attest to Roman and Byzantine agricultural influence on the spread of fruit crops such as peach, pear, plum, jujuba, apricot, cherry, pistachio nut, pine nut, and hazelnut, among others, and to Abbasid introduction of aubergines in the southern Levant.

Altogether, this evidence suggests that RAD was a greater force in the agricultural history of the first millennium CE than the IGR, which is also the current consensus from Iberia [39]. The significance of RAD is evident in the archaeobotany of additional regions, such as Italy, northwest Europe and Britain [34,38,68]. However, we should not dismiss the IGR on these grounds alone, since several of the proposed IGR crops are less likely to leave identifiable macroscopic traces (e.g., sugar cane, colocasia), and there is textual evidence for Early Islamic crop diffusion and agricultural development [22]. Hence it may be appropriate and productive to consider RAD and IGR part of the same process of first millennium CE agricultural development, as indicated by Early Islamic expansion of Roman and Byzantine crop introductions. Clearly the first millennium CE was an unprecedented period of change in local crop-plant species diversity in the eastern Mediterranean and beyond. The multi-regional evidence suggests that the multi-empire combination of Roman-Byzantine and Umayyad-Abassid regimes was a major force for crop diffusion, with a likely role for developments in the Sassanid empire underrepresented in current research. Yet the evidence presented here demonstrates that even the combined forces underlying first millennium CE crop diffusion affected, but did not immediately transform, people’s diets. At least until the end of that millennium, inhabitants of the Levant and Mediterranean region continued to rely primarily on long tried and tested Neolithic founder crops and early fruit domesticates.

Indeed, this situation widely persisted until the latter second millennium CE.

The new microregional data presented above supports an emerging multi-regional picture of both an unprecedented period for plant migrations and food diversity in the first millennium CE as well as gradual and incomplete local adoption. This is evident from Late Antique Negev Highlands archaeobotanical assemblages within which plants first attested to in the southern Levant during this period account for one third of the domesticated plant species diversity – more than any other period represented in the assemblage. Among these crops, only the aubergine represents an Early Islamic introduction, suggesting that Roman Agricultural Diffusion (RAD) was a greater force for intercontinental movement of crop plants than the proposed Islamic Green Revolution (IGR). However, both RAD and IGR plant species are very rare in the Negev Highlands assemblages, indicating slow incorporation into local foodways and agriculture. These findings present a window to a wider perspective on the last 11.5 millennia of southwest Asian crop diffusion, in which the first millennium CE is unprecedented for the diversity of plant species in motion yet consistent with a long-term pattern of gradual local adoption.

## Materials and Methods

Eleven middens from the three sites, Elusa, Shivta and Nessana, were excavated at approximately 10 cm spits to ensure chronological control. An intensive sampling-and-sifting strategy was followed to ensure optimal retrieval of plant remains (see **Supplementary Information**). Fine-sifted samples (see **Supplementary Information**) were sorted using an Olympus SZX9 stereo microscope and analyzed in the Bar-Ilan University Archaeobotany Lab. Course sifted samples were sorted by volunteers and archaeology students during the excavation and thereafter. Seed finds from the course sifting were examined and rare specimens taken to the Bar-Ilan University Archaeobotany Lab for identification. All identifications were made with reference to the Israel National Collection of Plant Seeds and Fruits at Bar-Ilan University. To confirm identification, the jujuba (*Ziziphus jujuba* Mill.) endocarp was scanned using a Bruker SkyScan 1174 desktop micro-CT scanner (**Supplementary Videos 1-2**). Identification criteria for this and other select specimens appear in the **Supplementary Information**. Information on previous archaeobotanical records of cultivated species was retrieved from the cited literature and lab records, as well as from online databases of archaeobotanical finds [71-73]. For palynological analysis, sediment samples from the middens were collected. However, all samples showed pollen barrenness, probably because of oxidation. Pollen from the reservoir and the northern church at Shivta did contribute additional taxa, as did wood and charcoal analyses. Results of pollen and wood analyses published by Langgut et al. [43,59] are summarized in **Supplementary Tables 5-6**.

The excavations’ stratigraphic, ceramic, and radiocarbon analyses enabled differentiation of five chronological phases obtained from the middens [43,54]: Roman (ca. 0–300 CE), Early Byzantine (ca. 300–450 CE), Middle Byzantine (ca. 450–550 CE), Late Byzantine (ca. 550– 650 CE) and Umayyad (ca. 650–750 CE), which was adjusted slightly based on radiocarbon dates presented herein. This enabled detection of trends within the Byzantine period as well as broader chronological comparisons. These periods are each represented by between one and four middens, and some middens span two periods (see **Table 2**). Grouping the seed/fruit crop remains into broad periods of introduction to the southern Levant was used to provide a general sketch of crop diffusion’s local influence in time.

## Data Availability

Only securely identified plant taxa are reported in the results of this study. All relevant data are included in the manuscript and supplementary materials. The investigated plant remains are currently stored in the Israel National Collection of Plant Seeds and Fruits at Bar-Ilan University and may be accessed by request to the authors.

## Acknowledgements

As part of a Ph.D. dissertation conducted at Bar-Ilan University, this research was supported by the Bar-Ilan Doctoral Fellowships of Excellence Program, the Rottenstreich Fellowship of the Israel Council for Higher Education, and the Molcho fund for agricultural research in the Negev. As part of the NEGEVBYZ project, this research was also supported by the European Research Council under the European Union’s Horizon 2020 Research and Innovation Programme (grant 648427) and the Israel Science Foundation (grant 340-14). Manuscript preparation was further supported by a Newton International Fellowship of the British Academy and a Marie S. Curie International Fellowship of the European Commission’s Horizon 2020 Framework Programme. Archaeology was conducted on behalf of the Zinman Institute of Archaeology, University of Haifa, under licenses of the Israel Antiquities Authority (Elusa: G-69/2014, G-10/2015, G-6/2017; Shivta: G-87/2015, G-4/2016; Nessana: G-4/2017). We also wish to thank the Israel Nature and Parks Authority for facilitating the excavations at Elusa, Shivta, and Nessana, as well as Ami and Dina Oach of Shivta Farm. For assistance with processing during the excavations, we are grateful to Ifat Shapira, Uri Yehuda, Ruti Roche, Gabriel Fuks, University of Haifa graduate students Aehab Asad, Ari Levy, and Yaniv Sfez, and countless other volunteers. We also wish to thank Y. Mahler-Slasky, Tammy Friedman, A. Hartmann-Shenkman, Michal David, Suembikya Frumin, I. Berko, and O. Bashari for laboratory assistance; Senthil Ram Prabhu Thangadurai and Prof. Ron Shahar of the Hebrew University of Jerusalem’s Laboratory of Bone Biomechanics for micro-CT scanning; and Sapir Haad for graphics.

## Supplementary Tables

**Supplementary Table 1.** Proposed IGR crops (according to Watson 1983 [16])

| Category | Latin name | English common name |
| --- | --- | --- |
| cereal | *Sorghum bicolor* (L.) Moench. | sorghum |
| cereal | *Oryza sativa* L. | rice |
| cereal | *Triticum durum* Desf. | hard wheat |
| textile | *Gossypium arboreum/herbaceum* L. | Old World cotton |
| tree fruit | *Citrus aurantium* L. | sour orange |
| tree fruit | *Citrus limon* L. | lemon |
| tree fruit | *Citrus aurantifolia* Swing. | lime |
| tree fruit | *Citrus grandis* L. | shaddock |
| tree fruit | *Musa sapientum/paradisiaca* L. | banana/plantain |
| tree fruit | *Cocos nucifera* L. | coconut |
| tree fruit | *Mangifera indica* L. | mango |
| vegetable | *Citrullus lanatus* (Thumb.) Mansf. | watermelon |
| vegetable | *Spinacia oleracea* L. | spinach |
| vegetable | *Cynara cardunculus* L. var. *scolymus* | artichoke |
| vegetable | *Colocasia antiquorum* Schott. | colocasia |
| vegetable | *Solanum melongena* L. | eggplant |
| condiment | *Saccharum officinarum* L. | sugar cane |

**Supplementary Table 2.** Proposed RAD crops (see main text for discussion and sources)

| Category | Latin name | English common name |
| --- | --- | --- |
| cereal | *Oryza sativa* L. | rice |
| cereal | *Sorghum bicolor* (L.) Moench. | sorghum |
| legume | *Lupinus albus* L. | white lupine |
| textile | *Cannabis sativa* L. | cannabis |
| tree fruit/nut | *Ceratonia siliqua* L. | carob |
| tree fruit/nut | *Morus nigra* L. | black mulberry |
| tree fruit/nut | *Prunus persica* (L.) Batsch | peach |
| tree fruit/nut | *Pyrus communis* L. | pear |
| tree fruit/nut | *Prunus domestica* L. | plum |
| tree fruit/nut | *Prunus armeniaca* L. | apricot |
| tree fruit/nut | *Prunus avium/cerasus* | cherry |
| tree fruit/nut | *Pistacia vera* L. | pistachio nut |
| tree fruit/nut | *Pinus pinea* L. | pine nut |
| tree fruit/nut | *Corylus avellana* L. | hazelnut |
| tree fruit/nut | *Ziziphus jujube* Mill. | jujuba |
| tree fruit/nut | *Citrus x limon* (L.) Osbeck | lemon |
| tree fruit/nut | *Cocos nucifera* L. | coconut |
| vegetable | *Cucumis melo* convar. *melo* | muskmelon |

**Supplementary Table 3.** Pre-1^st^ mill. CE Eastern Mediterranean introductions/domestications

| Period | Category | Latin name | English common name |
| --- | --- | --- | --- |
| Neolithic | cereal | *Triticum monococcum* L. subsp. *monococcum* | einkorn wheat |
| Neolithic | cereal | *T. turgidum* L. subsp. *dicoccum* (Schrank) Thell. | emmer wheat |
| Neolithic | cereal | *Hordeum vulgare* subsp. *vulgare* | barley |
| Neolithic | cereal | *Lens culinaris* L. | lentil |
| Neolithic | legume | *Pisum sativum* L. | pea |
| Neolithic | legume | *Cicer arietinum* L. subsp. *arietinum* | chickpea |
| Neolithic | legume | *Vicia ervilia* (L.) Willd. | bitter vetch |
| Neolithic | legume | *Vicia faba* L. | fava bean |
| Neolithic | fiber/oil | *Linum usitatissimum* L. | flax |
| Chalcolithic-Early Bronze | tree fruit/nut | *Olea europaea* L. | olive |
| Chalcolithic-Early Bronze | tree fruit/nut | *Vitis vinifera* L. | grapevine |
| Chalcolithic-Early Bronze | tree fruit/nut | *Ficus carica* L. | fig |
| Chalcolithic-Early Bronze | tree fruit/nut | *Ficus sycomorus* L. | sycomore |
| Chalcolithic-Early Bronze | tree fruit/nut | *Phoenix dactylifera* L. | date |
| Chalcolithic-Early Bronze | tree fruit/nut | *Punica granatum* L. | pomegranate |
| Chalcolithic-Early Bronze | tree fruit/nut | *Prunus amygdalus* Batsch. | almond |
| Bronze-Iron Age | tree fruit/nut | *Juglans regia* L. | walnut |
| Bronze-Iron Age | tree fruit/nut | *Citrus medica* L. | citron |
| Bronze-Iron Age | cereal | *Panicum miliaceum* L. | broomcorn millet |
| Bronze-Iron Age | cereal | *Setaria italica* (L.) P. Beauv. | foxtail millet |
| Bronze-Iron Age | legume | *Lathyrus clymenum* L. | Spanish vetchling |
| Bronze-Iron Age | legume | *Lathyrus sativus/cicera* L. | grass pea |
| Bronze-Iron Age | legume | *Trigonella foenum-graecum* L. | fenugreek |
| Bronze-Iron Age | condiment/oil | *Papaver somniferum* L. | opium poppy |
| Bronze-Iron Age | condiment/oil | *Nigella sativa* L. | black cumin |
| Bronze-Iron Age | condiment/oil | *Sesamum indicum* L. | sesame |
| Bronze-Iron Age | vegetable | *Citrullus lanatus* (Thunb.) Matsum. & Nakai | watermelon |

*Based primarily on Zohary et al. 2012 [3], this list includes only species whose evidence for domestication/introduction is* *clear. This and the preceding Supplementary tables are not meant as exhaustive lists but rather as a basis against which the* *Negev Highlands crop plant assemblage can be compared.*

**Supplementary Table 4.**
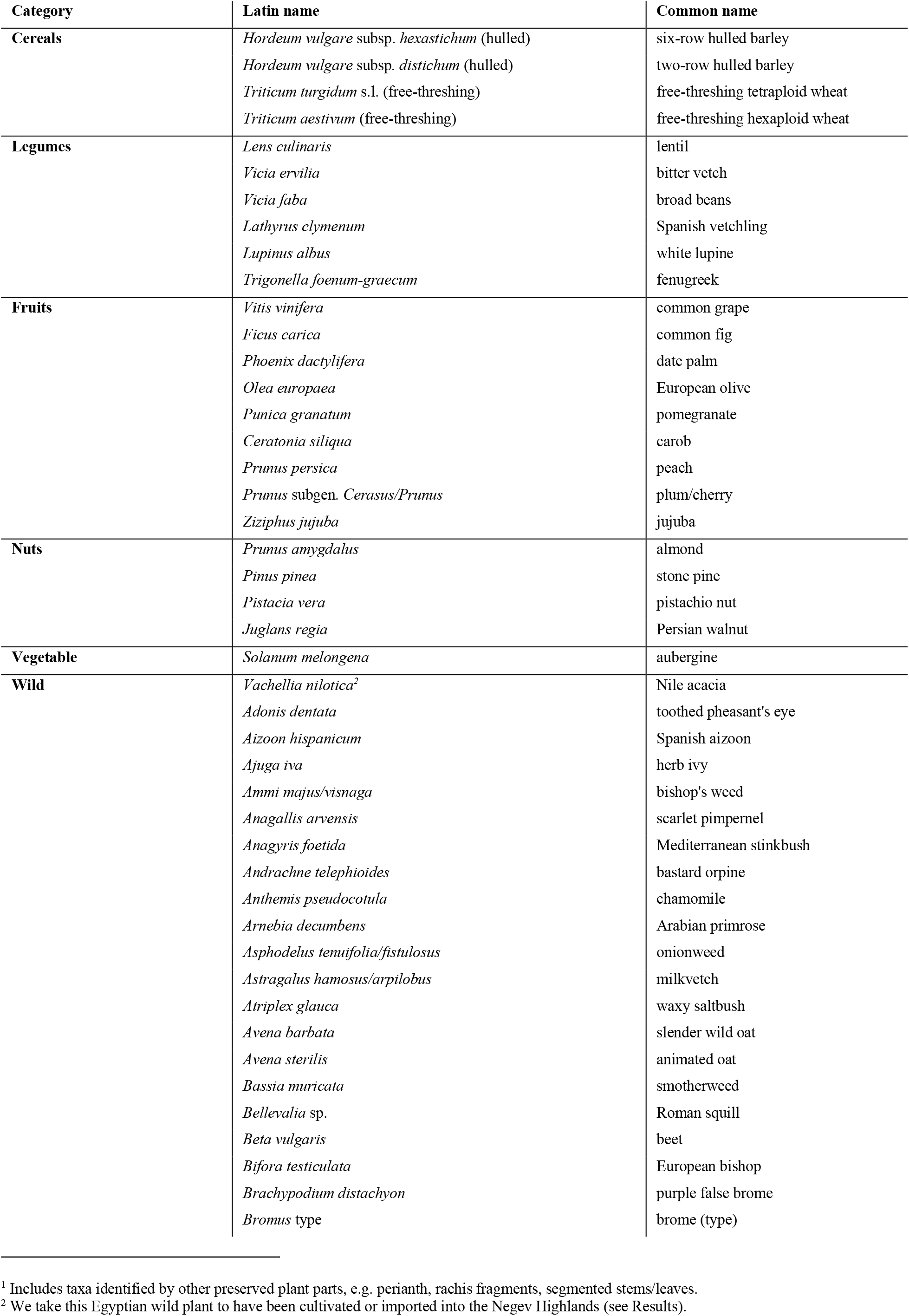
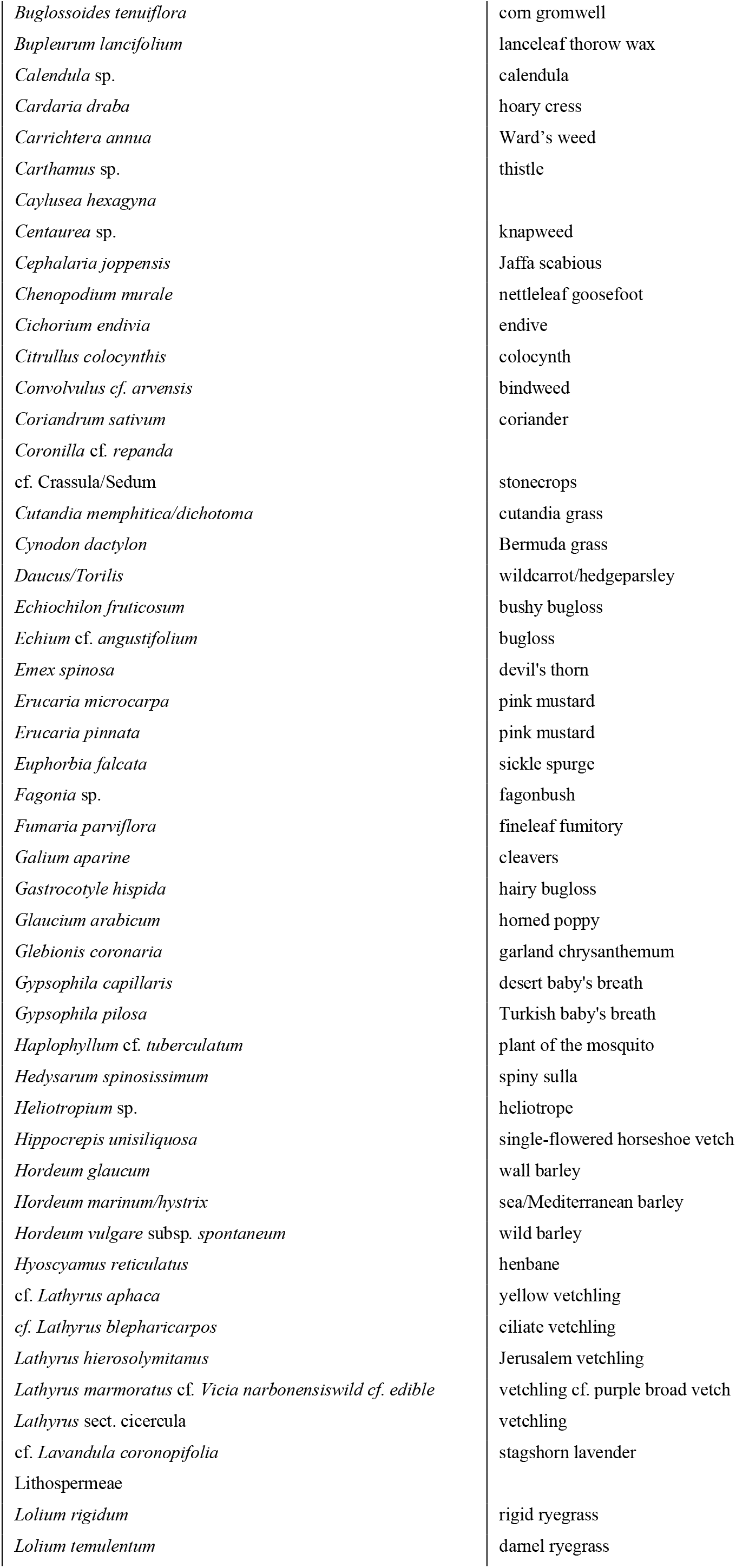
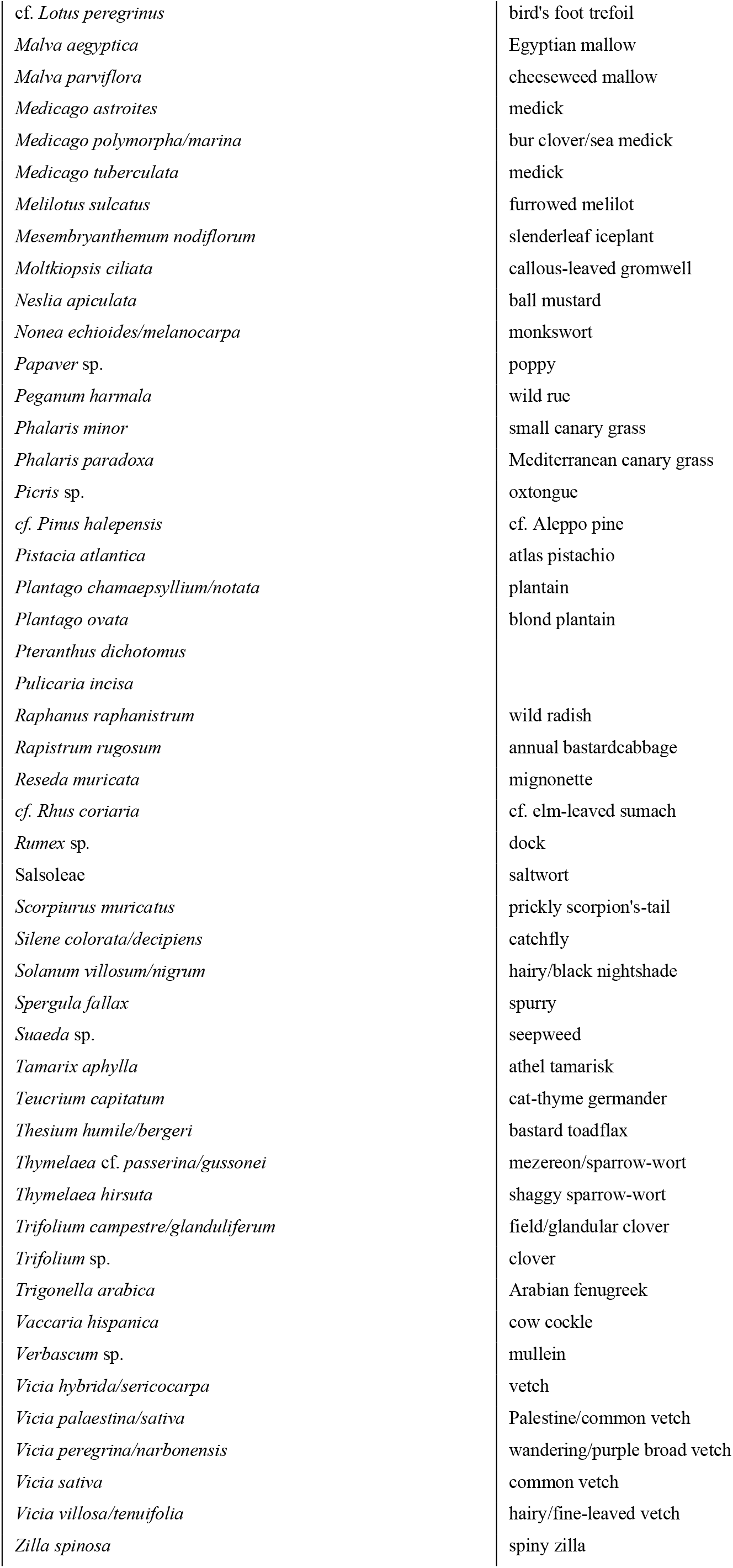
Carpological^1^ plant remains from Negev Highland middens

**Supplementary Table 5.** Identified wood and charcoal taxa from Shivta, Nessana and Elusa

| Category | Taxon | English common name | SVT | NZN | HLZ |
| --- | --- | --- | --- | --- | --- |
| Fruit trees | *Ficus carica* | common fig | + | + | + |
|  | *Ficus sycomorus* | Sycomore fig | - | + | + |
|  | *Hyphaene thebaica* | doum palm | + | + | - |
|  | *Olea europaea* | olive | + | - | + |
|  | *Phoenix dactylifera* | date palm | + | - | - |
|  | *Prunus* spp. (*dulcis/armeniaca*) | plum/apricot | + | + | - |
|  | *Punica granatum* | pomegranate | - | + | - |
|  | *Vitis vinifera* | grapevine | + | + | - |
| Exotic trees | *Buxus sempervirens* | boxwood | + | + | - |
|  | *Cedrus libani* | cedar of Lebanon | + | + | - |
|  | *Fraxinus excelsior* | European ash | - | + | - |
| Desert trees and shrubs | *Calotropis procera* | apple of Sodom | + | + | - |
|  | *Capparis spinosa* | caper bush | + | - | - |
|  | *Fagonia mollis* | fagonia | - | + | - |
|  | *Juniperus phoenicea* | Phoenician juniper | + | + | - |
|  | *Lycium* spp. | boxthorn | + | + | + |
|  | *Moringa peregrina* | Ben tree | + | - | - |
|  | *Pistacia atlantica* | Persian turpentine | + | + | - |
|  | *Populus/Salix* | poplar/willow | - | + | - |
|  | *Retama raetam* | white broom | + | + | + |
|  | *Rhamnus* spp. | buckthorn | + | + | + |
|  | *Salsola tetrandra* | saltwort [tetrandra] | + | - | - |
|  | *Salsola vermiculata* | Mediterranean saltwort | + | + | - |
|  | *Tamarix* spp. | tamarisk | + | + | + |
|  | *Ziziphus/Paliurus* | jujube/Jerusalem thorn | + | + | + |
|  | *Zygophyllum dumosum* | bushy bean caper | + | + | - |
| Mediterranean trees and shrubs | *Crataegus* spp. | hawthorn group/Maloideae | + | + | + |
|  | *Cupressus sempervirens* | Italian cypress | + | + | + |
|  | *Myrtus communis* | true myrtle | - | + | - |
|  | *Pinus halepensis* | Aleppo pine | + | + | + |
|  | *Pistacia palaestina* | terebinth | + | + | + |
|  | *Platanus orientalis* | oriental plane | - | + | + |
|  | *Quercus calliprinos* | Kermes oak | + | - | + |
|  | *Vitex agnus-castus* | chaste tree | - | + | - |

*Data for Shivta and Nessana derive from Langgut et al. 2021, Table 1 [59]; Data for Elusa are based* *on Bar-Oz et al. 2019, Table S8 [43].*

**Supplementary Table 6.**
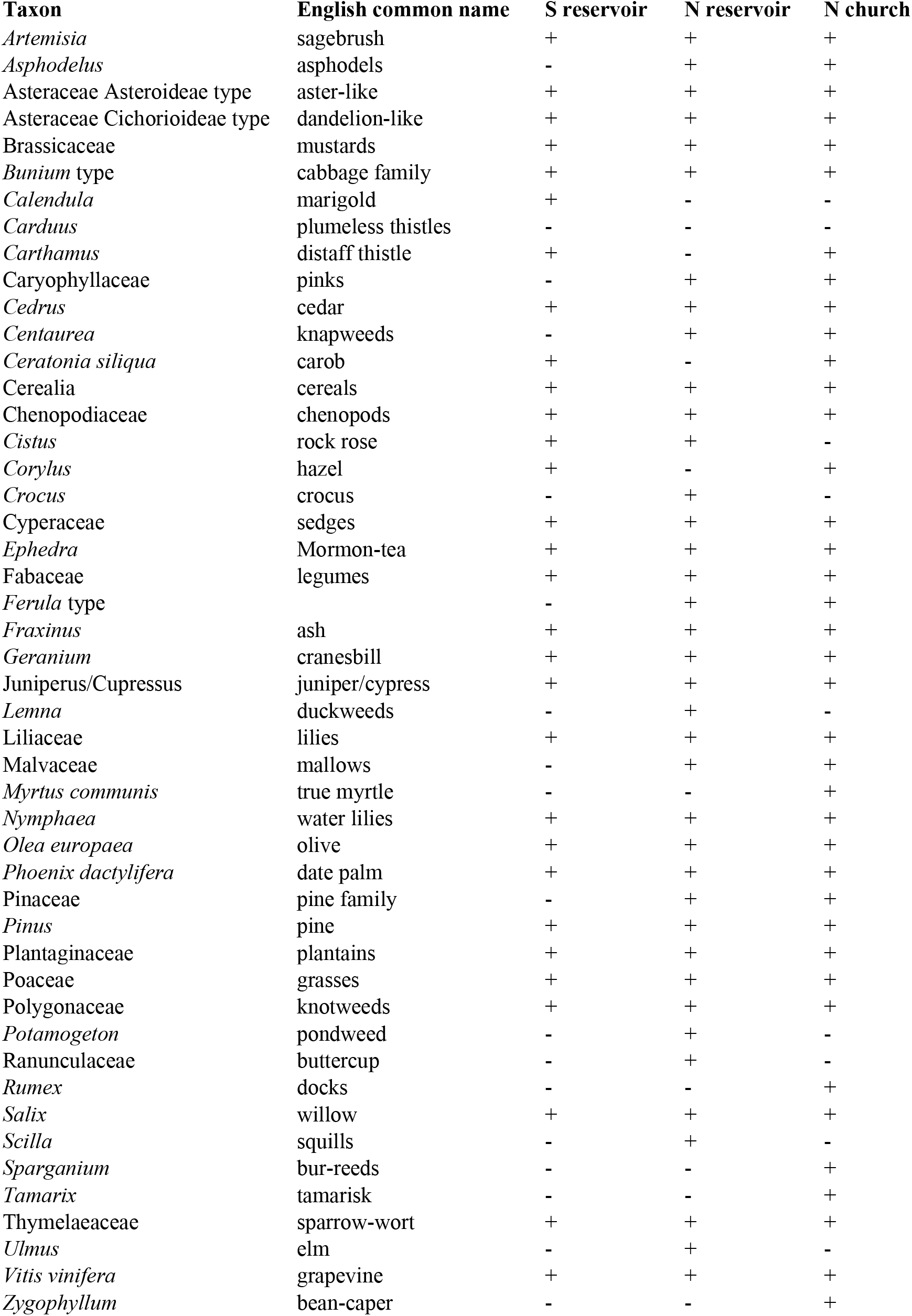
Identified pollen from Shivta reservoirs and garden

## Supplementary Information

### Field and laboratory extraction methods

Eleven middens from the three sites, Elusa, Shivta and Nessana, were excavated at approximately 10 cm height intervals to ensure chronological control (Figure 1 of main text). Loci and baskets were assigned by a combination of stratigraphy and sediment features during excavation. A three-pronged sifting strategy was adopted to maximize retrieval of artifacts and biological remains, while enabling complementary resolutions of analysis. All excavated material was sifted at one of three different levels, corresponding to sieve sizes: (1) Most excavated sediment was dry screened on site through 5 mm sieves. (2) Wet screening through 1 mm mesh was performed on two buckets (∼20 l) from each excavated locus-basket.

(3) One additional bucket from each locus-basket was set aside for fine screening. Selected buckets of sample sediments were divided into 3-liter subsamples which were processed by flotation or fine-mesh dry screening, and sieved using graduated sieves at 4 mm, 2 mm, 1 mm, 0.5 mm and sometimes 0.3 mm mesh sizes. One additional source of identified seeds was an assemblage of dissected charred dung pellets from two of the middens (Dunseth et al. 2019).

For ease of reference, (1) and (2) above are collectively referred to as *course sift samples* and (3) is referred to as *fine sift samples*. Due to the high volume of samples and the extremely high concentration of seeds within them, a subsampling strategy based on sieve mesh size was adopted for the fine sift samples. All flotation light fraction and heavy residues were sorted at the ≥ 2 mm mesh size. Light fraction was studied at 1 mm and 0.5 mm mesh sizes for select samples, such that at least three 1 mm samples and one 0.5 mm sample were sorted for each period on each site. Fine sift samples were sorted using an Olympus SZX9 stereo microscope. Course sifted samples were sorted by volunteers and archaeology students during the excavation and thereafter. Seed finds from the course sifting were visually examined with aid of a stereo microscope, and rare specimens taken to the Bar-Ilan University Archaeobotany Lab for identification.

On-site screening through 5mm sieves enabled very large volumes of sediment to be screened – nearly all excavated sediments were sifted in this way. As a result, course sifting demonstrated the ubiquity of dates and olives in all sites and periods, which would have been missed from fine sifting only. It also allowed for the discovery of less common large-seeded species; cherry/plum, pistachio, walnut, jujuba, fava bean and white lupine would have been missed entirely by exclusive fine sifting with its smaller sample volumes. This is reflected in the shorter species list in Table 2 of the main text, which records fine-sift retrieval only, in comparison with Table 1, which records course sift and fine sift retrieval. Since the same positive bias for retrieval of large seeds by 5mm sieves applies to both olive pits and date stones on one hand and those of cherry/plum, pistachio, walnut and jujuba on the other, this level of sifting facilitated the distinction between staple fruit crops and luxury/supplementary ones.

Wet screening through 1 mm mesh also allowed for processing of a greater sample volume (up to 20 l per locus-basket) than for the fine sift samples (3 l per locus-basket), providing additional qualitative and quantitative data for most of the major domesticated plant seeds. Ratios of cereal grains to grape pips from wet screening and fine sifting were shown to be equivalent, enabling wet-screened samples to complement fine-sifted samples in quantitative analysis (Fuks et al. 2020). Wet screening through 1 mm mesh and sorting by volunteers is a cost-effective method for discovering the main domesticated plant species on site, but it provides incomplete coverage.

As long-recognized in archaeobotany, fine-mesh sifting enabled retrieval of a much wider range of plants. Without it, we would have entirely missed the presence of fig drupelets on site, let alone their high ubiquity. Evidence for crop processing, especially of cereals, derived exclusively from the fine sifting, as did the vast majority of wild/weed seeds. In addition, the subsampling strategy by mesh size proved highly effective in maximizing species retrieval and quantitative comparison between contexts. Sorting 100% of fine sift sediments at the 2 mm+ mesh size enabled full recovery of all major domesticated species except figs.

Subsampling material retrieved from 1 mm and 0.5 mm sieves enabled a balance to be met between constraints and coverage of small finds. These sieve sizes produced the bulk of cereal rachis fragments, fig drupelets and remains of most identified wild/weed taxa.

Altogether, the above multi-pronged sifting strategy effectively maximized retrieval of plant remains and contributed to the high diversity of identified taxa. This, together with the focus on organically rich rubbish middens and a multi-site micro-regional approach produced a dataset that is relevant on a macro-regional and Holocene-wide scale.

### Seed identification

Identifications were performed with reference to the Israel National Collection of Plant Seeds and Fruits at Bar-Ilan University. Cereal grain morphometry was employed to identify candidates, using the Computerized Key of Grass Grains developed by Mordechai Kislev’s laboratory (Kislev et al. 1992; 1997; 1999). As aids to identification and analysis, local plant guides were consulted, particularly the *Flora Palaestina* (Zohary and Feinbrun-Dothan, 1966–1986). Additional floras of Mediterranean, Irano-Turanian and Saharo-Arabian phytogeographic regions were consulted as needed (Townsend and Guest 1966–1985; Meikle, 1977, 1985; Zohary et al. 1980–1994; Feinbrun-Dothan et al. 1998; Turland, 1993; Boulos, 1999-2005; Davis, 1966–2001; Danin, 2004). To confirm identification, the jujuba (*Ziziphus jujuba*) endocarp was scanned using a micro-CT (Bruker desktop SkyScan 1174) at the Laboratory of Bone Biomechanics, Hebrew University of Jerusalem (Supplementary Videos 1-2).

Identification criteria for rare, domesticated plant specimens discussed in the main text are summarized below:

### Aubergine (Solanum melongena L.)

*S. melongena* and other *Solanum* seeds are laterally compressed, broadly oval-shaped and under 5 mm in maximal length. *S. melongena* seeds are distinguished from wild *Solanum* seeds of the southern Levant by their larger size, reticulated seed coat pattern, and the wide ovoid hilum set in a recess in the seed’s lateral outline (Van der Veen and Morales 2011: 93; Amichay and Weiss 2020]: 679). This includes *S. incanum* L. which was identified at Byzantine Ein Gedi and is considered by some to be the wild progenitor of *S. melongena* (Melamed and Kislev 2005). The latter two criteria also distinguish *S. melongena* from domesticated *Capsicum* spp. Based on these criteria, we identified three definitive *S. melongena* seeds from Umayyad Shivta (Area E, Locus 504, Basket 5029). Poor preservation precludes definitive identification for an additional three fragmented seeds from Umayyad Nessana (Locus 102) for which *S. melongena* nonetheless appears to be the only candidate (SI Figure 1).

**SI Figure 1.**
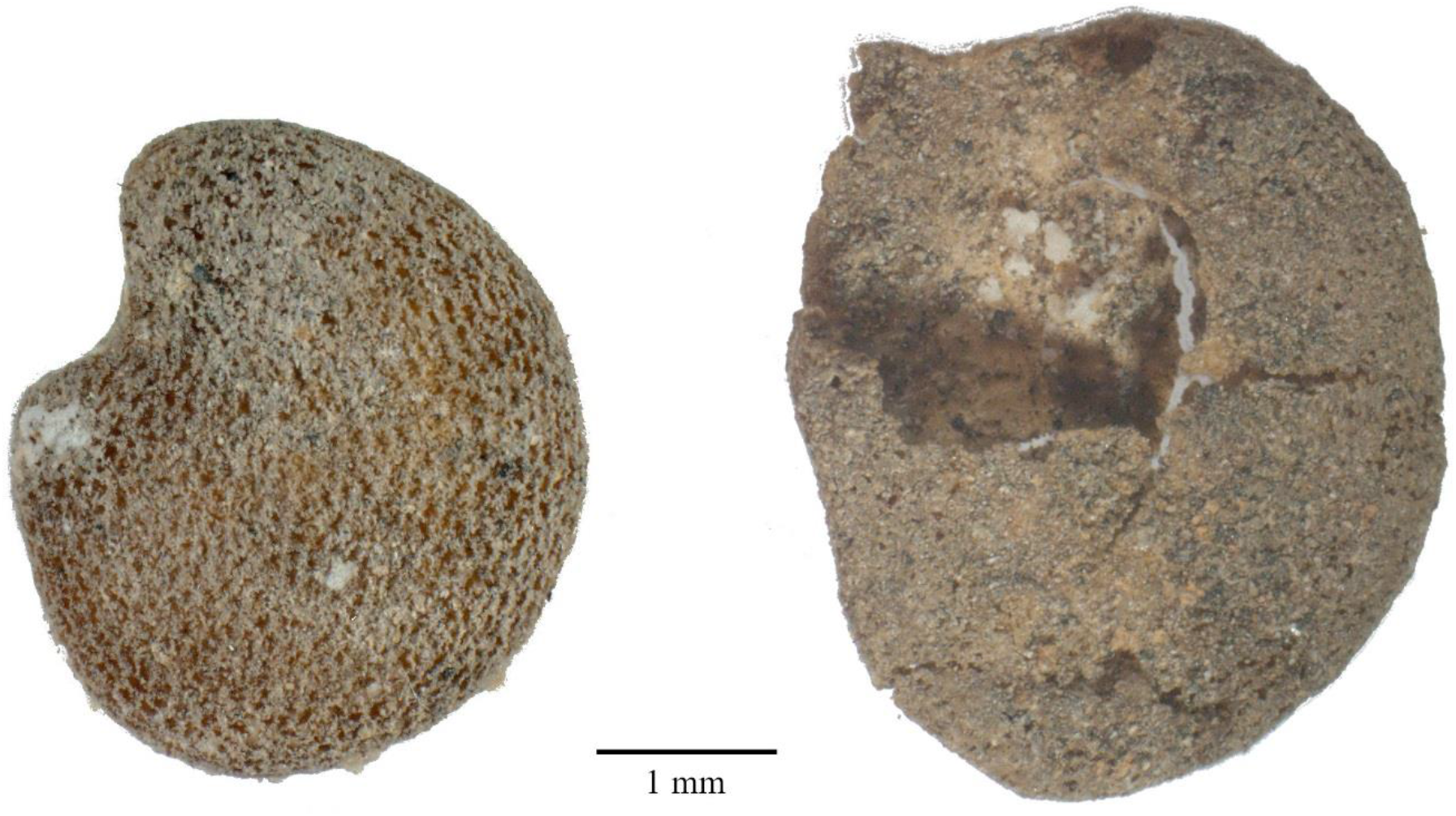
Left: *Solanum melongena* L. seed from Shivta (E 504-5029). Right: cf. *Solanum melongena* from Nessana (A 102-1072-1).

### Cherry/plum (Prunus subgen. Cerasus/Prunus)

A single ovoid endocarp with a pointed apex, elliptical base (5 mm by 2.5 mm), and smooth surface was found in a course-sift sample from Umayyad Shivta (Area K1, Locus 165, Basket 1652; SI Figure 2). Its length from apex to base is 12.67 mm, width 9.33 mm, and breadth

7.67 mm. A ventral ridge runs down the length of the endocarp, from apex to base, accompanied by two ridges on either side and at equal distance from the central ridge. However, the right ventral ridge exists only on the top third of the endocarp while the left ventral ridge is visible in the top two thirds. The dorsal side is marked by a single longitudinal ridge. The above characteristics ruled out apricot, peach, and almond, and leave cherry and plum as candidates (*Prunus* subgen. *Cerasus/Prunus*). Due to the wide variety of plum and cherry cultivars (Depypere et al. 2007) not fully covered by the reference collection, we did not identify to species.

**SI Figure 2.**
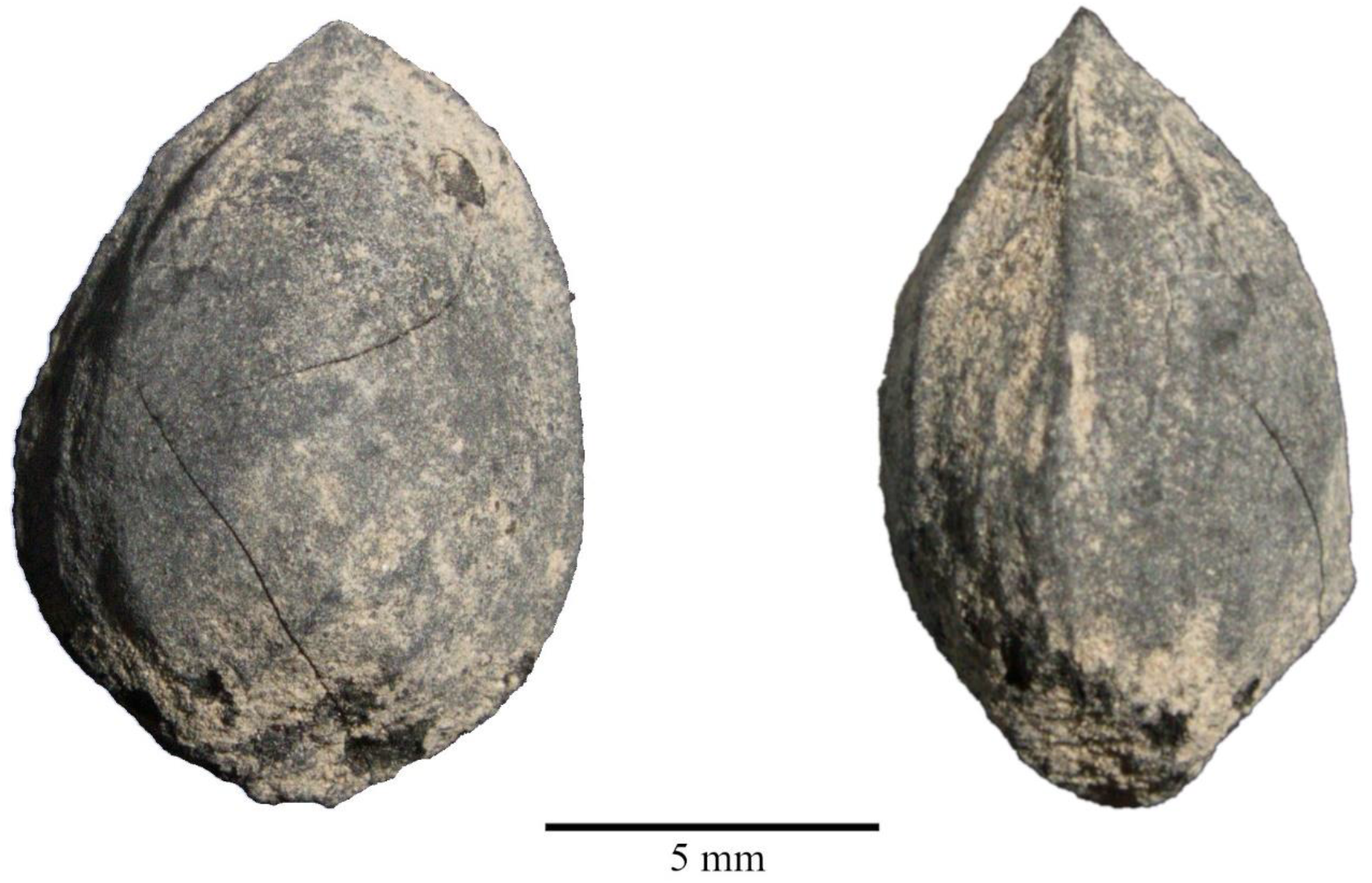
*Prunus* subgen. Cerasus/Prunus endocarp from Shivta (K1 165-1652)

### Jujuba (Ziziphus jujuba Mill.)

A single charred obconical-mucronate endocarp was found from Umayyad-period layers from Shivta (Area E, Locus 501, Basket 5108). Micro-CT scanning (using a Bruker desktop SkyScan 1174), demonstrated it to be spherically hollow with remnants of a partition (see Supplementary Videos 1-2), confirming its status as a fruit endocarp. The external endocarp dimensions (11.16 mm x 6.0 mm x 5.33 mm) and obconical with markedly narrowing apex (SI Figure 3) are unique to certain varieties of *Ziziphus jujuba*. The specimen’s pointed edges tapered slightly and the external grooves characteristic of *Z. jujuba* are barely recognizable, apparently the result of abrasion during or following charring. Remnants of the characteristic v-shaped basal scar between the two endocarp halves (Jiang et al. 2013, their Fig. 6) are barely visible, again likely due to abrasion. Species with similar endocarps include local wild types of *Ziziphus* (Z. *spina-christi*, Z. *lotus*, Z. *nummalaria*), however these are always spherical and never obconical-mucronate to the extent of *Z. jujuba* and the specimen at hand.

**SI Figure 3.**
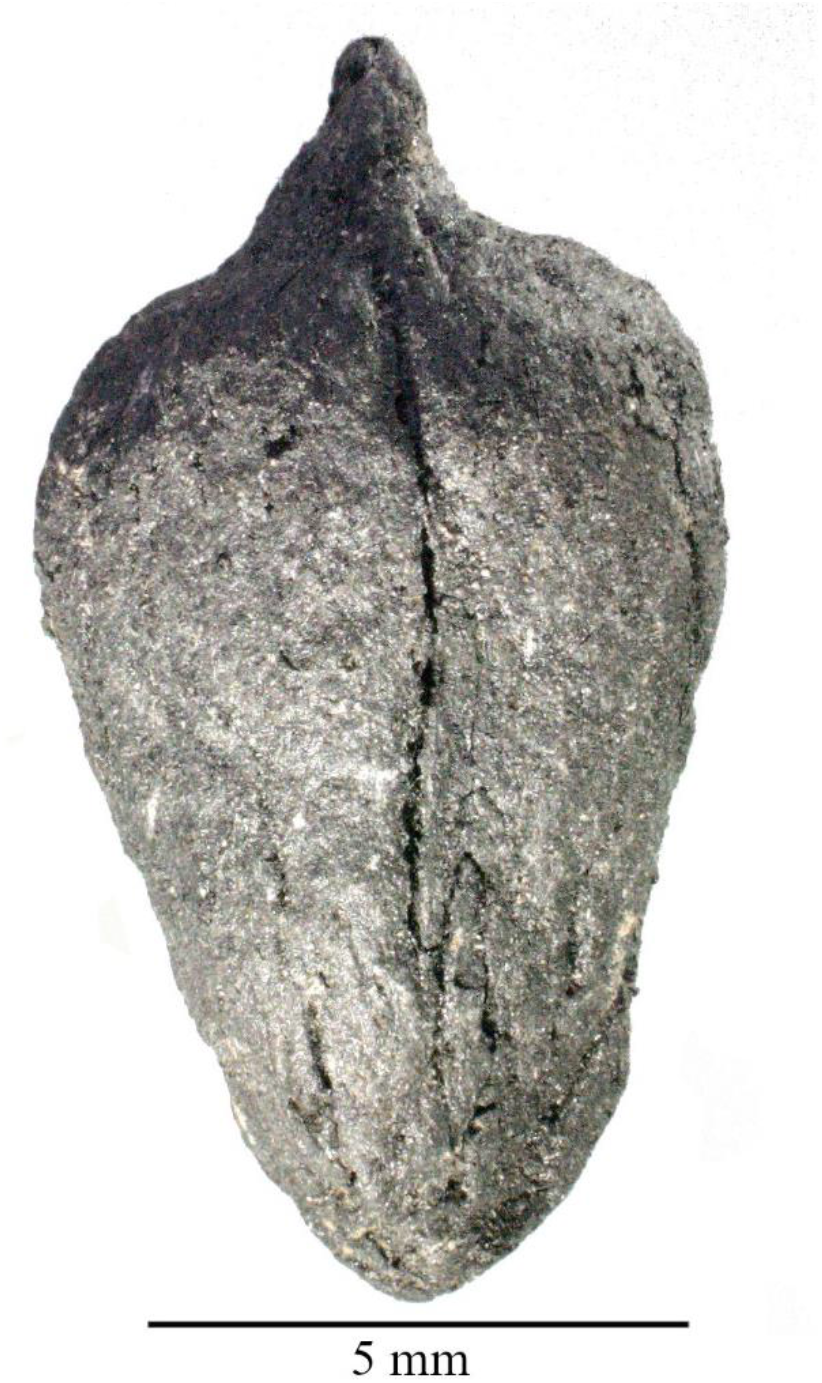
*Ziziphus jujuba* Mill. endocarp from Shivta (E 501-5108)

### Nile acacia (Vachellia nilotica (L.) P.J.H.Hurter & Mabb.)

*Vachellia* (syn. *Acacia*) is a genus in the Mimosoideae subfamily of the Fabaceae. Seeds of Mimosoideae species native to the southern Levant are elliptical to ovate and compressed. On each face of the seedcoat a conspicuous pleurogram delimits an ovate areole (Gunn 1984; Al-Gohary and Mohamed 2007). The pleurogram may either be open-ended, i.e. U-shaped/horseshoe-shaped, or closed, concentric to the seed contour. To identify seeds with these traits found in the middens, we compared seeds of Mimosoideae species native to the southern Levant, based on samples in the Israel National Collection of Plant Seeds and Fruits:

i. *Vachellia nilotica* (L.) P.J.H.Hurter & Mabb.) syn. *Acacia nilotica* (L.) Willd. ex Delile;
ii. *Senegalia laeta* (R.Br. ex Benth.) Seigler & Ebinger syn. *Acacia laeta* R.Br. ex Benth.;
iii. *Acacia pachyceras* O. Schwartz; (iv) *Vachellia tortilis* subsp. *raddiana* (Savi) Kyal. & Boatwr. syn. *Acacia raddiana* Savi; (v) *Vachellia tortilis* (Forssk.) Galasso & Banfi syn. *Acacia tortilis* (Forssk.) Hayne; (vi) *Faidherbia albida* (Delile) A.Chev.; and (vii) *Prosopis farcta* (Banks & Sol.) J.F.Macbr. We observed that *V. nilotica* seeds are distinguished by the following characteristics:

1. The pleurogram’s border (linea fissura) is closed, creating an ovate areole (SI Figure 4).
2. The areole is largest, relative to seed size, in *V. nilotica*, i.e., the distance from the linea fissura to the seed edge is shortest in this species (SI Table 1).
3. The areole’s widest part is in the top third of the seed (SI Table 1; SI Figure 4).
4. A protrusion is present next to the hilum which we observed to be unique to *V. nilotica* seeds among the above species.

*V. nilotica* seeds tend to be the largest of the above except for *P. farcta*, although interspecies diversity leads to size overlap between *V. nilotica, A. pachyceras* and *V. tortilis* subsp. *raddiana* (SI Table 1). *P. farcta* seeds are like *Vachellia* spp. seeds in shape but tend to be larger than most *Vachellia* seeds and more ovate to pear-shaped. Their pleurograms are visibly open. *V. nilotica* seeds were identified using a combination of criteria (1)-(4) above in midden samples from Elusa (Area A1, Locus 1/10a; A4, L. 4/06a-4/07a; SI Figure 4). Remains of *Vachellia* were identified also in other Negev Highland sites: One seed from Nessana (A, L. 125, B. 1446) was identified as *Vachellia* sp., while a single seed from Shivta (K1, L. 153, B. 1579) could only be identified as *Vachellia*/*Prosopis farcta* due to poor preservation.

**SI Figure 4.**
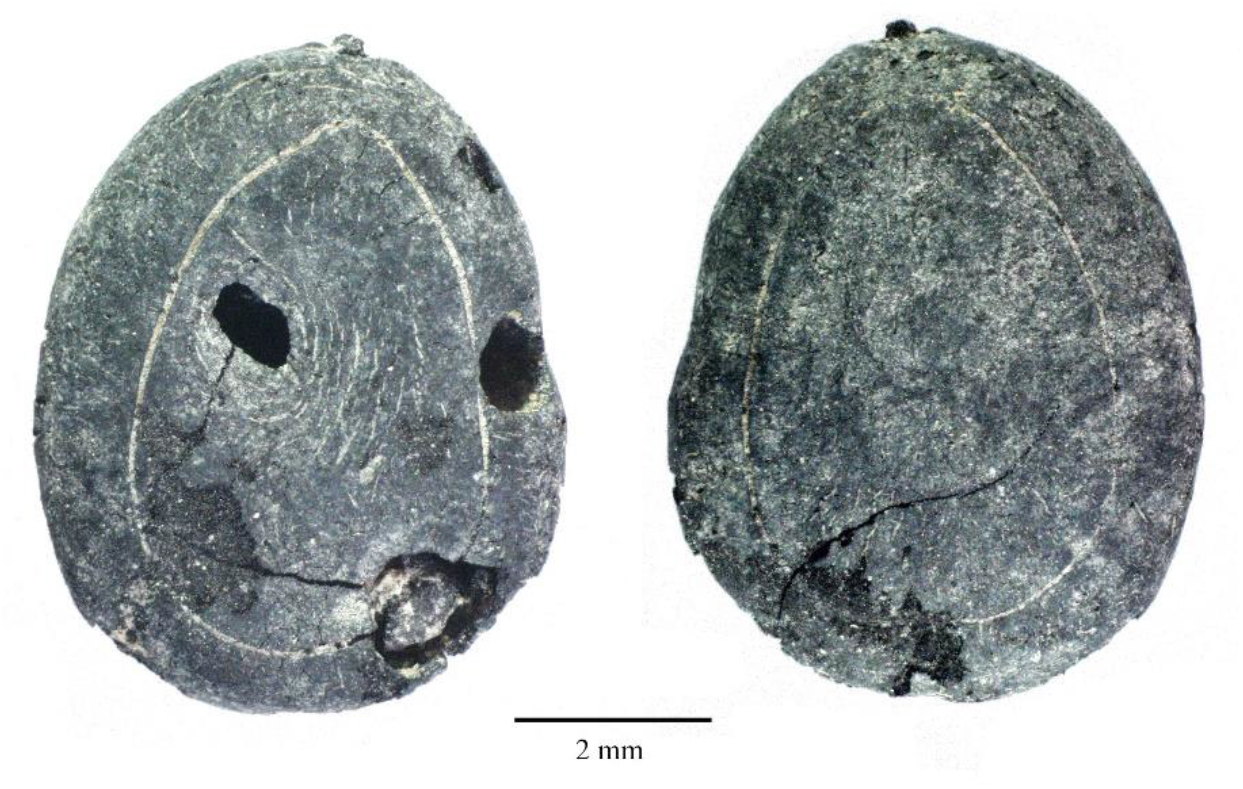
*Vachellia nilotica* (L.) Willd. ex Delile seed faces A and B from Elusa (A1/10a)

**SI Table 1.** *Some* Acacia *spp. seed measurements from the Israel National Collection of Plant Seeds and Fruits*

| Species | Population | seed<br># | seed<br>face<br>(A/B) | seed<br>length<br>(mm) | seed<br>width<br>(mm) | areole<br>length<br>(mm) | areole<br>width<br>(mm) | (seed width-<br>areole width)/<br>seed width | (seed length-<br>areole length)/<br>seed length | max.<br>areole<br>width |
| --- | --- | --- | --- | --- | --- | --- | --- | --- | --- | --- |
| *A. nilotica* | Elusa A, archaeological | 1 | A | 7.5 | 6.0 | 6.0 | 4.2 | 0.30 | 0.20 | a |
| *A. nilotica* | Elusa A, archaeological | 1 | B | 7.5 | 6.0 | 6.1 | 4.1 | 0.32 | 0.19 | a |
| *A. nilotica* | Elusa B, archaeological | 2 | A | 5.7 | 4.7 | 5.3 | 3.6 | 0.23 | 0.07 | a |
| *A. nilotica* | Luxor 1981 | 3 | A | 10.0 | 7.6 | 9.0 | 6.0 | 0.21 | 0.10 | a |
| *A. nilotica* | Luxor 1981 | 3 | B | 10.1 | 7.7 | 8.9 | 5.6 | 0.27 | 0.12 | a |
| *A. nilotica* | Luxor 1981 | 4 | A | 10.5 | 7.7 | 8.9 | 6.0 | 0.22 | 0.15 | a |
| *A. nilotica* | Luxor 1981 | 4 | B | 10.5 | 7.7 | 8.8 | 6.0 | 0.22 | 0.16 | a |
| *A. nilotica* | Luxor 1981 | 5 | A | 10.9 | 7.3 | 9.5 | 5.7 | 0.22 | 0.13 | a |
| *A. nilotica* | Luxor 1981 | 5 | B | 10.6 | 7.0 | 9.5 | 5.0 | 0.29 | 0.10 | a |
| *A. nilotica* | Luxor 1981 | 6 | A | 7.0 | 6.5 | 6.2 | 4.5 | 0.31 | 0.11 | a |
| *A. nilotica* | Luxor 1981 | 6 | B | 7.0 | 6.4 | 6.0 | 4.5 | 0.30 | 0.14 | a |
| *A. pachyceras* | Wadi Ram 26.2.95 | 7 | A | 9.2 | 6.9 | 6.2 | 3.2 | 0.54 | 0.33 | c |
| *A. pachyceras* | Wadi Ram 26.2.95 | 7 | B | 9.1 | 6.7 | 6.6 | 3.2 | 0.52 | 0.27 | c |
| *A. pachyceras* | Wadi Ram 26.2.95 | 8 | A | 10.5 | 8.0 | 7.4 | 4.2 | 0.48 | 0.30 | a |
| *A. pachyceras* | Wadi Ram 26.2.95 | 8 | B | 10.5 | 8.0 | 7.8 | 3.9 | 0.51 | 0.26 | a |
| *A. pachyceras* | Wadi Ram 26.2.95 | 9 | A | 10.6 | 6.5 | 7.9 | 3.5 | 0.46 | 0.25 | b |
| *A. pachyceras* | Wadi Ram 26.2.95 | 9 | B | 10.4 | 6.4 | 7.8 | 3.6 | 0.44 | 0.25 | b |
| *A. pachyceras* | Nahal Hayyun 15.3.71 | 10 | A | 8.1 | 5.7 | 5.0 | 2.7 | 0.53 | 0.38 | a |
| *A. pachyceras* | Nahal Hayyun 15.3.72 | 10 | B | 8.0 | 5.7 | 5.1 | 2.5 | 0.56 | 0.36 | a |
| *A. pachyceras* | Nahal Hayyun 15.3.73 | 11 | A | 8.5 | 5.8 | 6.4 | 3.3 | 0.43 | 0.25 | b |
| *A. pachyceras* | Nahal Hayyun 15.3.74 | 11 | B | 8.5 | 5.8 | 6.3 | 3.2 | 0.45 | 0.26 | b |
| *A. pachyceras* | Nahal Hayyun 15.3.75 | 12 | A | 7.7 | 6.2 | 6.0 | 3.6 | 0.42 | 0.22 | b |
| *A. pachyceras* | Nahal Hayyun 15.3.76 | 12 | B | 7.5 | 6.2 | 6.0 | 3.6 | 0.42 | 0.20 | b |
| *A. raddiana* | Moje Awad | 13 | A | 7.9 | 5.8 | 5.4 | 3.5 | 0.40 | 0.32 | e |
| *A. raddiana* | Moje Awad | 13 | B | 7.9 | 5.7 | 5.2 | 3.5 | 0.39 | 0.34 | e |
| *A. raddiana* | Moje Awad | 14 | A | 9.7 | 6.5 | 7.0 | 3.8 | 0.42 | 0.28 | d |
| *A. raddiana* | Moje Awad | 14 | B | 9.6 | 6.5 | 7.0 | 4.0 | 0.38 | 0.27 | c |
| *A. raddiana* | Moje Awad | 15 | A | 8.1 | 5.9 | 5.5 | 3.8 | 0.36 | 0.32 | c |
| *A. raddiana* | Moje Awad | 15 | B | 8.0 | 6.0 | 5.7 | 3.5 | 0.42 | 0.29 | c |
| *A. raddiana* | Ein Gedi 19.5.1917 | 16 | A | 8.0 | 5.5 | 5.9 | 3.5 | 0.36 | 0.26 | c |
| *A. raddiana* | Ein Gedi 19.5.1917 | 16 | B | 8.0 | 5.5 | 5.6 | 3.4 | 0.38 | 0.30 | c |
| *A. raddiana* | Ein Gedi 19.5.1917 | 17 | A | 8.0 | 5.3 | 5.9 | 3.5 | 0.34 | 0.26 | c |
| *A. raddiana* | Ein Gedi 19.5.1917 | 17 | B | 8.1 | 5.4 | 5.5 | 3.4 | 0.37 | 0.32 | c |
| *A. raddiana* | Ein Gedi 19.5.1917 | 18 | A | 8.0 | 5.4 | 5.9 | 3.5 | 0.35 | 0.26 | c |
| *A. raddiana* | Ein Gedi 19.5.1917 | 18 | B | 8.0 | 5.4 | 5.8 | 3.4 | 0.37 | 0.28 | c |

Table uses *Acacia* as used in the reference accessions; for synonyms see text above. Max. areole width is based on distance from hilum: a ) upper third (from hilum); b) upper third-midway; c) midway; d) midway-lower third; e) lower third

### Spanish vetchling (Lathyrus clymenum L.)

Identification of *Lathyrus clymenum* was based on morphological similarity to ancient *L. clymenum* seeds identified from Tel Nami by Kislev (1993). Diagrams and measurements reported by Sarpaki and Jones (1990) for a large number of *L. clymenum* seeds from Late Bronze Age Akrotiri and Knossos were also used.

The following generalized description refers to the identified *L. clymenum* seeds from Shivta and Nessana: The seeds are laterally compressed, nearly rectangular circumstance. In lateral view, the radicle lies on the short side, perpendicular to the long side where the hilum lies (SI Figure 5). The radicle forms a somewhat planar face, especially by comparison with the other sides of the seed. The dorsal side (parallel to that on which the hilum lies), is conspicuously carinated, whereas the ventral side was only moderately carinated. The hilum occupies over half the length of the ventral side. It begins at one end of the ventral side (near the radicle) and ends just before the circular lens. The thin seed coat is neither perfectly smooth nor tuberculate but appears grainy at magnification of ca. 40X.

*L. clymenum* seeds were identified at Nessana, midden A (106-1255 cf. 106-1257; 101-1032) and several from midden K at Shivta (153-1588,1610; 158-1618; 166-1658; 169-1678,1703; 172-1689). The positions, shapes and relative sizes of the hilum and lens matched those of the Tel Nami *L. clymenum* seeds and the depictions in Sarpaki and Jones (1990). The same is true for seed coat thickness and texture, as well as the markedly carinated dorsal side. One seed from Shivta (K1, 153-1588) measured below than the range of Tel Nami seed dimensions (SI Table 2). However, its relative dimensions and clear morphology justified unequivocal identification as *L. clymenum*.

**SI Table 2.** Select L. clymenum seed measurements from Tel Nami

| Seed | L (mm) | B (mm) | T (mm) | L/B | L/T |
| --- | --- | --- | --- | --- | --- |
| 1 | 4.3 | 2.3 | 3.6 | 1.87 | 1.19 |
| 2 | 4.6 | 2.4 | 3.9 | 1.92 | 1.18 |
| 3 | 4.2 | 2.2 | 3.05 | 1.91 | 1.38 |
| 4 | 3.6 | 2.75 | 2.9 | 1.31 | 1.24 |
| 5 | 3.3 | 2.5 | 3.5 | 1.32 | 0.94 |
| mean | 4.00 | 2.43 | 3.39 | 1.66 | 1.19 |
| s.d. | 0.48 | 0.19 | 0.37 | 0.29 | 0.14 |

**SI Figure 5.**
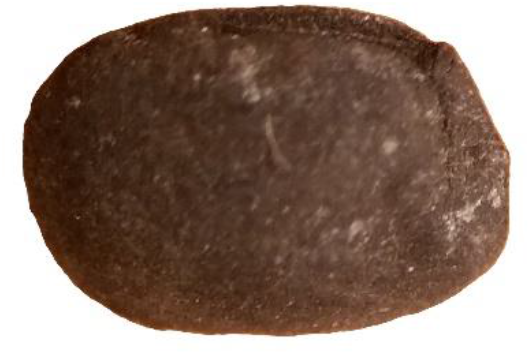
*Lathyrus clymenum* L. seed from Shivta, midden K. Length ca. 3.5 mm.

### White lupine (Lupinus albus L.)

Three species of lupine (*Lupinus*) which grow today in the southern Levant are distinct for their large (ca. 1 cm), compressed quadrangular seeds: *L. palaestinus, L. pilosus*, and the cultivated *L. albus*. Viewed laterally, the seeds of these species have a near-circular, or D-shaped outline and, frequently, a visible depression or dimple. The triangular radicle forms the perimeter’s straightest side, while the hilum leads from the radicle tip toward the lens at an angle such that the lens and radicle are on perpendicular sides with the hilum cutting across between the two. The lens is nearly as large as the hilum and both are elliptic. The seed coat surrounds the hilum by a characteristic elliptical protrusion. Throughout, the seed coat consists of at least two layers visibly distinct in cross-section, with the outer layer having a smooth surface and the inner layer having a grainy surface. As is common among domesticated legumes in general, the seed coat of cultivated *L. albus* is much thinner than its local wild relatives. An additional feature distinguishing *L. albus* seeds from *L. palaestinus/pilosus* is the presence of a clear transverse ridge separating the radicle depression and the hilum on the seed surface. In *L. palaestinus/pilosus*, by contrast, the radicle depression and hilum are essentially contiguous, running smoothly one into the other.

Three candidates for lupine seeds were identified among course-sifted archaeobotanical remains from Nessana (Area A, Locus 101, Baskets 1008/1 and 1040/2). The single seed from Basket 1040 (SI Figure 6) is compressed with a lateral depression and a near-circular quadrangle in outline measuring 70 × 75 mm. Remains of a triangular radicle on the seed’s straight side are clearly visible. These features narrowed its identification to one of the three aforementioned *Lupinus* species. Both lens and hilum are visible; their shape and orientation match those of *Lupinus* seeds. A slight but clear protrusion separating the hilum from the radicle depression warrant identification as *Lupinus albus*. Remnants of a thin and grainy seed coat are visible in the center of the cotyleda’s surface, in the middle of the lateral depression.

Two additional seeds from Basket 1008/1 show characteristic lupine (*Lupinus* sp.) hila and radicle. The seeds measure 65 × 70 mm and 75 × 80 mm which, together with their D-shaped outlines, corresponds with that typical to the large lenticular lupine species mentioned above. The two seeds from basket 1008/1 are broader than the *L. albus* seed from Basket 1040/2, and the characteristic lateral depression is not visible. This is apparently due to lateral swelling and partial disfiguration during charring as is common in charred legume seeds. In the larger of the two seeds, a thin, grainy seed coat is visible surrounding the triangular radicle and covering one of the cotyleda. In that same seed, a topographic separation between the radicle depression and hilum justifies identification as *L. albus*.

**SI Figure 6.**
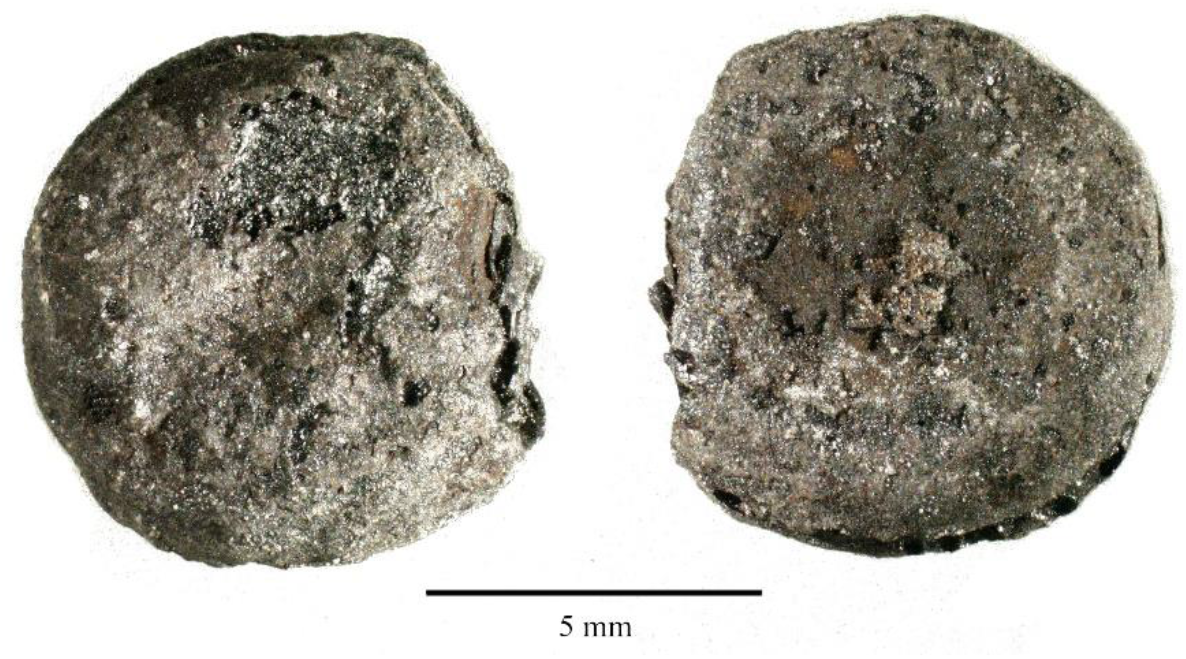
*Lupinus albus L*. seed faces A and B from Nessana (A 101-1040/2)

### Radiocarbon dating

Periodization of the studied assemblages followed those used by Fuks et al. (2020), based on ceramic typologies and previous radiocarbon dates (Bar-Oz et al. 2019). In this study we dated the loci-baskets containing unprecedented finds for southern Levantine archaeobotany, as well as the locus containing well-preserved aubergine seeds in Shivta. The aubergine, lupin and jujuba seeds were too rare to sacrifice for direct radiocarbon so barley grains were selected from the very same sediment sample within each locus-basket. Radiocarbon dating was performed by the Poznan Radiocarbon Laboratory, and calibration was made with the OxCal v4.4.2 (Bronk Ramsey 2020), using atmospheric data from Reimer et al (2020). All dates reflect assemblages from the Early Islamic period (Table 4).

Although the calibrated ranges vary, the sample containing aubergine (*S. melongena*) falls within the Abbasid period at the 95% confidence level; samples containing white lupin (*L. albus*) and jujuba (*Z. jujuba*) are either Umayyad or from the early Abbasid period (mid-7^th^ – late 8^th^ c. cal. CE).

### Micro-CT scanning

Micro-CT scans on the *Z. jujuba* endocarp were conducted by Senthil Ram Prabhu Thangadurai at the Laboratory of Bone Biomechanics, Hebrew University of Jerusalem. Optical resolution (pixel size): 9.6 µm; exposure: 4500 ms; rotation step: 0.400 degrees; 180 degree rotation option was used; 0.25 mm thick aluminium filter. The scans confirmed identification as an endocarp by revealing its hollow inner structure and partition. For full identification criteria see above. The following scanning files are attached to this article:

*SI Video 1* – Micro-CT longitudinal scans of *Z. jujuba* endocarp.

*SI Video 2* – Micro-CT lateral scans of *Z. jujuba* endocarp.

